# Multidimensional analysis of Gammaherpesvirus RNA expression reveals unexpected heterogeneity of gene expression

**DOI:** 10.1101/482042

**Authors:** Lauren M. Oko, Abigail K. Kimball, Rachael E. Kaspar, Ashley N. Knox, Carrie B. Coleman, Rosemary Rochford, Tim Chang, Benjamin Alderete, Linda F. van Dyk, Eric T. Clambey

**Author notes:** **Co-Corresponding authors:** Eric T. Clambey, Linda F. van Dyk. **Lead author** for MS correspondence.

## Abstract

Virus-host interactions are frequently studied in bulk cell populations, obscuring cell-to-cell variation. Here we investigate endogenous herpesvirus gene expression at the single-cell level, combining a sensitive and robust fluorescent in situ hybridization platform with multiparameter flow cytometry, to study the expression of gammaherpesvirus non-coding RNAs (ncRNAs) during lytic replication, latent infection and reactivation in vitro. This method allowed robust detection of viral ncRNAs of murine gammaherpesvirus 68 (γHV68), Kaposi’s sarcoma associated herpesvirus and Epstein-Barr virus, revealing variable expression at the single-cell level. By quantifying the inter-relationship of viral ncRNA, viral mRNA, viral protein and host mRNA regulation during γHV68 infection, we find heterogeneous and asynchronous gene expression during latency and reactivation, with reactivation from latency identified by a distinct gene expression profile within rare cells. Further, during lytic replication with γHV68, we find many cells have limited viral gene expression, with only a fraction of cells showing robust gene expression, dynamic RNA localization, and progressive infection. Lytic viral gene expression was enhanced in primary fibroblasts and by conditions associated with enhanced viral replication, with multiple subpopulations of cells present in even highly permissive infection conditions. These findings, powered by single-cell analysis integrated with automated clustering algorithms, suggest inefficient or abortive γHV infection in many cells, and identify substantial heterogeneity in viral gene expression at the single-cell level.

**AUTHOR SUMMARY:** The gammaherpesviruses are a group of DNA tumor viruses that establish lifelong infection. How these viruses infect and manipulate cells has frequently been studied in bulk populations of cells. While these studies have been incredibly insightful, there is limited understanding of how virus infection proceeds within a single cell. Here we present a new approach to quantify gammaherpesvirus gene expression at the single-cell level. This method allows us to detect cell-to-cell variation in the expression of virus non-coding RNAs, an important and understudied class of RNAs which do not encode for proteins. By examining multiple features of virus gene expression, this method further reveals significant variation in infection between cells across multiple stages of infection, even in conditions generally thought to be highly uniform. These studies emphasize that gammaherpesvirus infection can be surprisingly heterogeneous when viewed at the level of the individual cell. Because this approach can be broadly applied across diverse viruses, this study affords new opportunities to understand the complexity of virus infection within single cells.

## INTRODUCTION

The *Herpesviridae* are a family of large dsDNA viruses that include multiple prominent human and animal pathogens [1]. Although these viruses infect different cell types, and are associated with diverse pathologies, they share conserved genes and two fundamental phases of infection: lytic replication and latent infection [1]. Lytic replication is characterized by a cascade of viral gene expression, active viral DNA replication and the production of infectious virions. Conversely, latency is characterized by limited viral gene expression and the absence of de novo viral replication. While latent infection is a relatively quiescent form of infection, the herpesviruses can reactivate from latency, to reinitiate lytic replication.

Among the herpesviruses, the gammaherpesviruses (γHV) are lymphotropic viruses that include the human pathogens Epstein-Barr virus (EBV) [2] and Kaposi’s sarcoma associated herpesvirus (KSHV) [3]. Murine gammaherpesvirus 68 (γHV68, or MHV-68; ICTV nomenclature *Murid herpesvirus 4*, MuHV-4), is a well-described small animal model for the γHVs [4]. While these viruses establish a lifelong infection that is often clinically inapparent, immune-suppressed individuals are particularly at risk for γHV-associated malignancies [5].

Herpesvirus gene expression is extremely well-characterized in bulk populations. Despite increasing evidence for single-cell heterogeneity in gene expression [6-8], there remains limited understanding of herpesvirus infection at the single-cell level [9-12]. Here, we tracked endogenous viral and host RNAs using a sensitive, robust fluorescent in situ hybridization assay combined with multiparameter flow cytometry (PrimeFlow™) [13] to analyze the expression and inter-relationships of viral ncRNA, viral mRNA and cellular mRNA at the single-cell level during γHV latency, reactivation and lytic replication. These studies revealed unanticipated heterogeneity of infection, emphasizing how single-cell analysis of virus infection can afford significant new insights into the complexity of γHV infection.

## RESULTS

### Single-cell analysis of viral RNAs during lytic infection

Traditional measurements of gene expression frequently rely on pooled cellular material, obscuring intercellular variation in gene expression. To better define expression of γHV RNAs at the single cell level, we employed the PrimeFlow™ RNA assay [13] to study viral gene expression during murine gammaherpesvirus 68 (γHV68) infection, a small animal γHV [4, 13]. This method is a highly sensitive, extremely specific in situ hybridization assay, integrating Affymetrix-designed branched DNA technology with single-cell analysis powered by multiparameter flow cytometry. This method has been successfully used to detect both virus and host RNAs (e.g. in the context of HIV infected individuals [13, 14]).

We first tested the ability of PrimeFlow™ to measure multiple viral RNAs during lytic infection with γHV68, including small non-coding RNAs (<u>t</u>RNA-<u>m</u>iRNA <u>e</u>ncoding <u>R</u>NAs or TMERs [15]) and mRNAs. Mouse 3T12 fibroblasts were infected with a multiplicity of infection (MOI=5 plaque forming units (PFU) of virus/cell). Under these conditions, TMER-5, one of the eight γHV68 TMERs, and the γHV68 ORF73, were readily detectable by conventional real-time PCR in γHV68-infected, but not mock-infected, cultures (Fig. 1A, C). Parallel cultures were analyzed for RNA expression by PrimeFlow™. Whereas mock-infected cells had no detectable expression of either the γHV68 TMERs or ORF73, WT γHV68-infected fibroblasts had a prominent population of TMER+ and ORF73+ cells, respectively (Fig. 1B and 1D). Infection of cells with a TMER-deficient γHV68 (TMER-TKO [16]), in which TMER expression is ablated through promoter disruption, revealed no detectable TMER expression (Fig. 1B), yet robust ORF73 expression (Fig. 1D). Parallel studies revealed ready detection of ORF18, another γHV68 gene product (Fig. 1E). These studies show that PrimeFlow™ is a sensitive, robust and specific method to detect both viral non-coding and messenger RNAs during lytic infection, quantifying both the frequency of gene expression and expression on a per cell basis.

**Figure 1.**
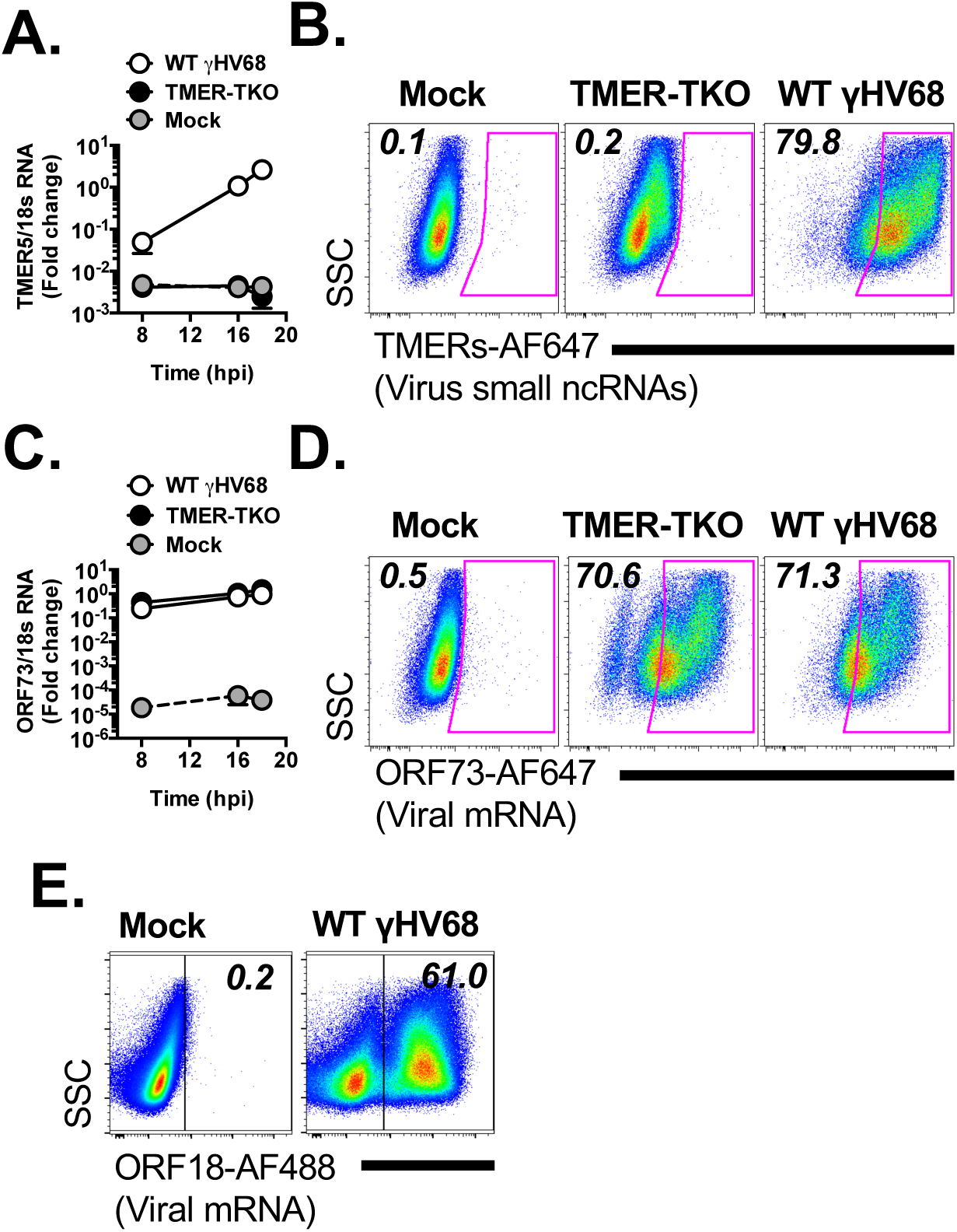
RNA-Flow cytometry using the PrimeFlow™ method affords robust and sensitive analysis of endogenous γHV genes at the single-cell level. Viral RNA analysis in γHV68-infected fibroblasts by qRT-PCR (A, C) or by flow cytometric analysis using PrimeFlow™ at 16 hpi (B, D, E, F). Samples were either mock, TMER-TKO, or WT γHV68-infected, with infections done using 5 plaque forming units/cell. qRT-PCR standardized to 18s RNA, at the indicated times. All flow cytometric events gated on a generous FSC x SSC gate, followed by singlet discrimination. PrimeFlow™ analysis quantified probe fluorescence for (B) TMERs, (D) ORF73, or (E) ORF18, relative to side-scatter area (SSC-A). Probe fluorescence is indicated, with all probes detected using either AlexaFluor (AF) 647 or 488 conjugates. Data representative of 2 independent experiments, each done with biological replicates.

### Heterogeneous gene expression during γHV latency and reactivation from latency

γHV latency is characterized by limited gene expression. We next measured viral RNAs during latency and reactivation using the γHV68-infected A20 HE2.1 cell line (A20.γHV68), a drug-selected latency model with restricted viral gene expression that can reactivate following stimulation [17]. A20.γHV68 cells are characterized by restricted viral gene expression, yet remain competent for reactivation from latency and the production of infectious virions following chemical stimulation with the phorbol ester, TPA [17, 18].

When we compared TMER expression between uninfected (parental, virus-negative A20) and infected (A20.γHV68) cells by qRT-PCR, the viral ncRNA TMER-5 was readily detectable in A20.γHV68 cells above background signals in parental A20 cells, with minimal changes between untreated and chemically-stimulated conditions (Fig. 2A). PrimeFlow™ analysis of TMER expression in untreated A20.γHV68 cells revealed that a majority of these cells expressed the TMERs, as defined by a positive signal in samples subjected to the TMER probe relative to unstained cells (Fig. 2B). Untreated A20.γHV68 cells contained a high frequency of cells expressing intermediate levels of TMERs (i.e. TMER^mid^ cells), with a significant signal enrichment above parental, virus-negative A20 cells (Fig. 2C). While the frequency of TMER^mid^ cells remained relatively constant following treatment with TPA (compare “Untreated” versus “TPA stimulated”, Fig. 2C), TPA stimulated A20.γHV68 cultures also contained a small fraction of cells with high levels of TMERs (i.e. TMER^high^ cells), not present in untreated cultures (Fig. 2C-D). Chemical stimulation is known to result in variable penetrance of reactivation in latently infected cell lines [17]. Based on this, we hypothesized that these rare, TMER^high^ cells may represent a subset of cells that are undergoing reactivation from latency.

**Figure 2.**
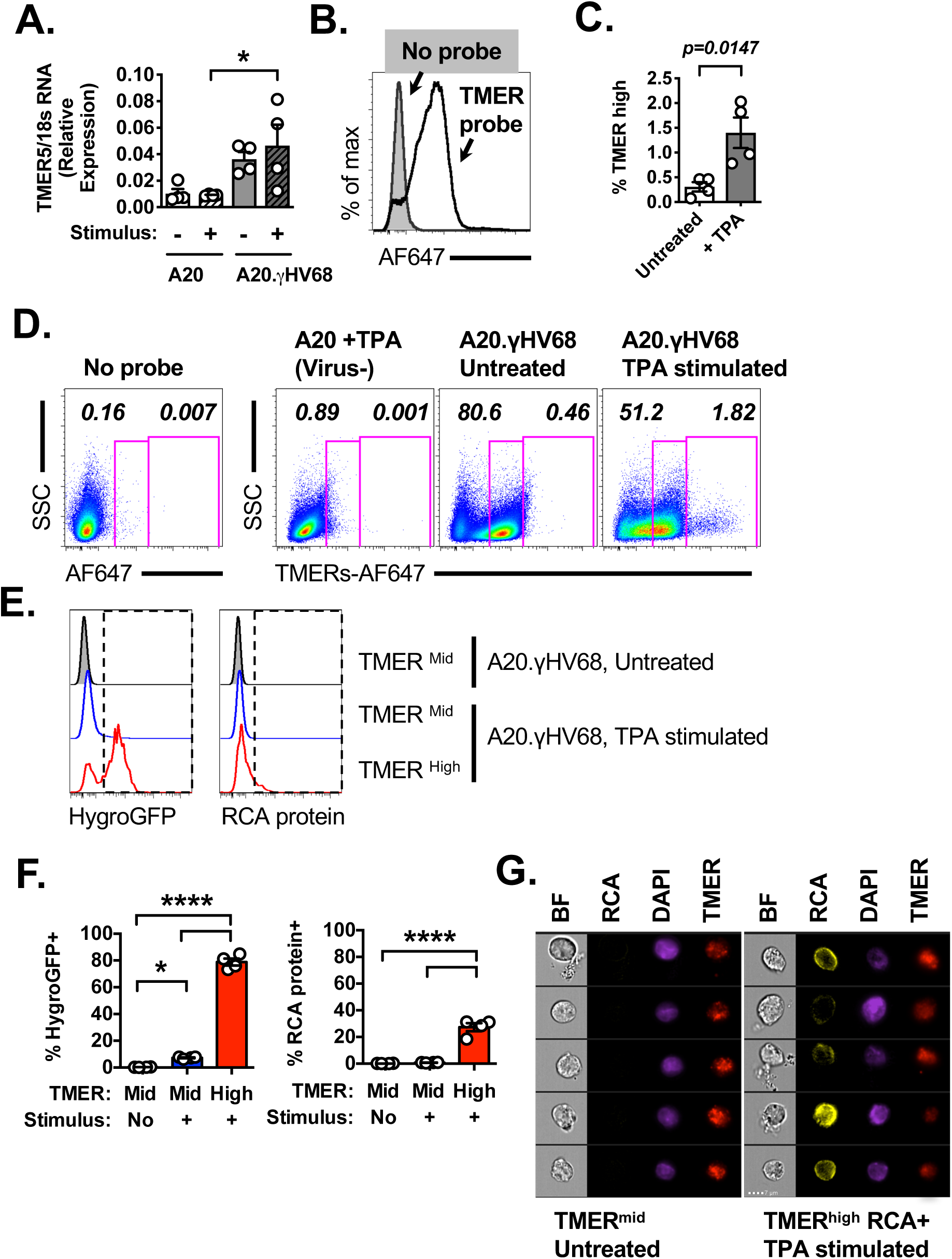
Heterogeneous gene expression in a γHV68 latently infected B cell line. Viral, host RNA analysis during γHV68 latency and reactivation in A20.γHV68 (HE2.1) cells by qRT-PCR (A) and flow cytometric analysis using PrimeFlow™ (B-G), comparing untreated or TPA-stimulated samples at 24 hrs post-treatment. Analysis includes A20, virus-negative cells and A20.γHV68 cells. (A) qRT-PCR analysis of TMER-5 expression relative to 18s RNA in A20 and A20.γHV68 (HE2.1) cells, untreated or stimulated with TPA for 24 hrs. (B) PrimeFlow™ detection of TMER expression in A20.γHV68 (HE2.1) cells, comparing either samples that were unstained (solid gray) or stained for the TMERs (open black line). (C) Analysis of TMER expression in multiple conditions, comparing untreated and stimulated A20 and A20.γHV68 cells, with gates defining the frequency of events that expressed either intermediate (mid) or high levels of TMERs. Data depict lymphocytes that were singlets, defined by sequential removal of doublets. (D) Quantification of the frequency of TMER^high^ cells in stimulated A20.γHV68 cells. (E) Histogram overlays of HygroGFP and RCA protein expression in A20.γHV68 cells comparing TMER^mid^ cells from untreated cultures (top, gray), TMER^mid^ cells from TPA-stimulated cultures (middle, blue), with TMER^high^ cells from stimulated cultures (bottom, red). (F) Quantification of the frequencies of HygroGFP+ (left) and RCA protein+ (right) cells as a function of TMER expression and treatment condition. (G) Flow cytometric analysis on an imaging flow cytometer, with each row showing an individual cell and representative images of brightfield (BF), RCA protein (RCA), DAPI, and TMER localization in TMER^mid^ cells from untreated cultures (left) or TMER^high^ RCA+ cells from stimulated (right) A20.γHV68 cells. Data are from two independent experiments, with biological replicates within each experiment for all A20.γHV68 cultures. Graphs depict the mean ± SEM, with each symbol identifying data from a single replicate. Statistical analysis was done using an unpaired t test (D) or one-way ANOVA, subjected to Tukey’s multiple comparison test (A, F), with statistically significant differences as indicated, * p<0.05, **** p<0.0001.

To test this, we analyzed the properties of TMER^mid^ and TMER^high^ cells, comparing viral protein expression in untreated and stimulated A20.γHV68 cells. We analyzed: 1) a γHV68 expressed GFP-hygromycin resistance fusion protein (HygroGFP), under the control of a heterologous viral promoter (the CMV immediate early promoter) [17], and 2) the γHV68 regulator of complement activation (RCA), a viral protein encoded by the γHV68 ORF4, an early-late transcript [19]. The vast majority of TMER^mid^ cells were negative for HygroGFP and RCA (i.e. HygroGFP- RCA-), regardless of whether the cells were present in untreated or stimulated cultures (Fig. 2E-F). Conversely, TMER^high^ cells, which were present at an increased frequency in stimulated cultures, had a significantly increased frequency of HygroGFP+ cells with induction of RCA protein+ cells in a subset of cells when compared to TMER^mid^ cells present in either untreated or stimulated cultures (Fig. 2E-F). By using imaging flow cytometry, we further analyzed the subcellular localization of TMERs in TMER^mid^ cells compared with TMER^high^ RCA+ cells. TMERs were predominantly nuclear in both TMER^mid^ and TMER^high^ RCA+ cells, as defined by co-localization with DAPI fluorescence (Fig. 2G). These data demonstrate that the TMERs are expressed during latency, and that following reactivation-inducing stimulation, TMERs are further induced in a rare subset of cells which are characterized by increased viral transcription and translation.

### Detection of endogenous viral gene expression during KSHV latency and reactivation

To extend these findings, we analyzed viral gene expression in the KSHV infected B cell tumor line, BCBL-1, focused on detection of an abundant viral ncRNA, the KSHV polyadenylated nuclear RNA (PAN, nut1, or T1.1) [20]. PAN RNA is known to be highly inducible upon induction of reactivation in KSHV latently infected B cell lymphoma cell lines [10, 20]. The frequency of PAN RNA+ cells was low in untreated BCBL-1 cells, with ~1% of cells spontaneously expressing this ncNRA (Fig. 3A-B). Despite the low frequency, this hybridization was clearly above background, as ncRNA defined on the KSHV- and EBV-negative B cell lymphoma cell line BL41 [21, 22] (Fig. 3A-B). Upon stimulation of BCBL-1 cells with the reactivation-inducing stimuli TPA and sodium butyrate, the frequency of PAN RNA+ cells significantly increased with expression in ~25% of cells (Fig. 3A-B). Although stimulation of BCBL-1 cells significantly increased the frequency of PAN RNA+ events compared to untreated cultures, PAN RNA expression on an individual cell basis was comparable between cells from untreated or stimulated cultures (Fig. 3C). As anticipated, stimulation of BCBL-1 cells was associated with increased viral DNA, consistent with stimulated cultures undergoing reactivation from latency (Supplemental Fig. 1).

**Figure 3.**
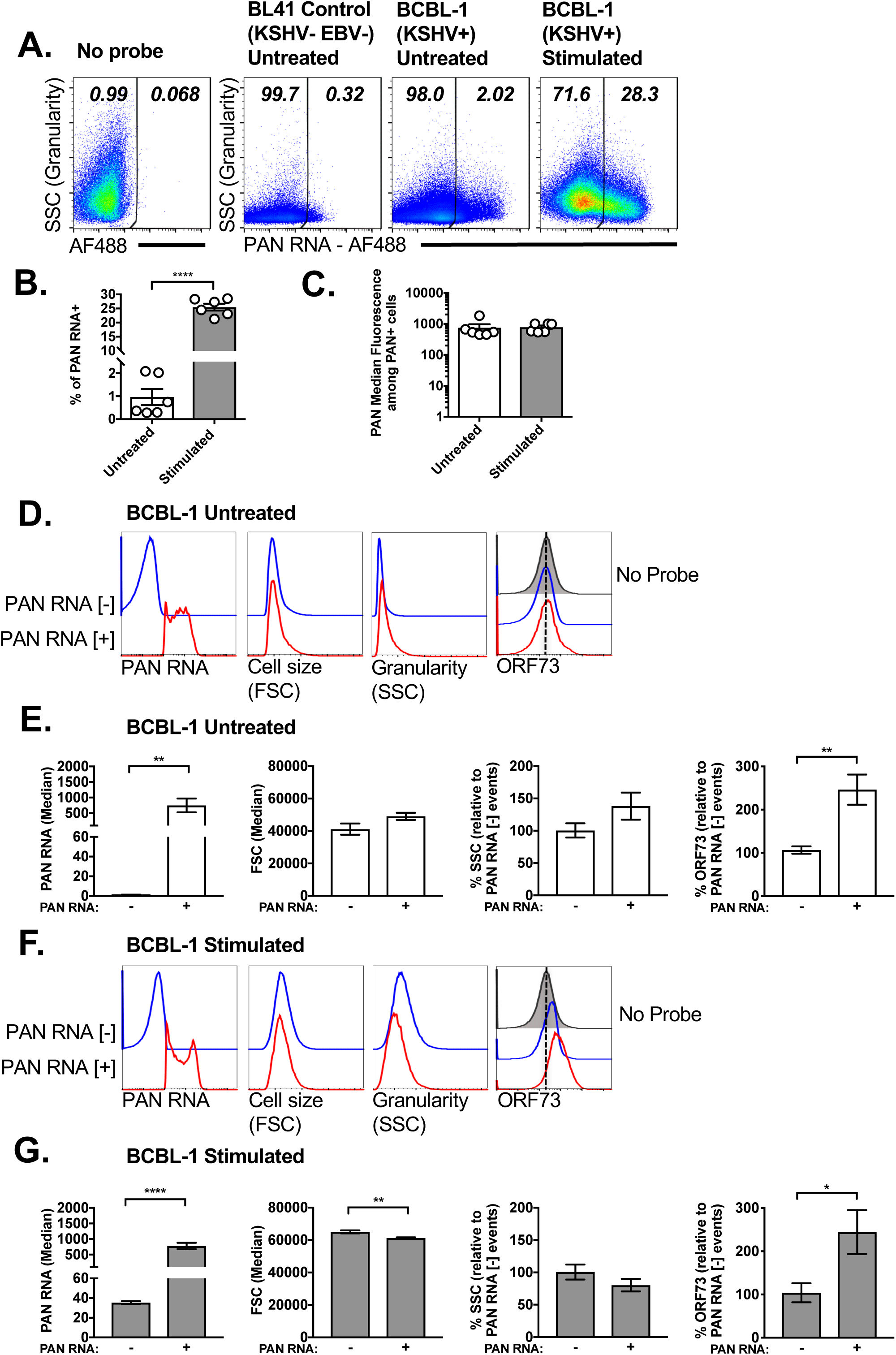
Single-cell analysis of KSHV PAN RNA expression in the BCBL-1 B cell lymphoma cell line. (A) Flow cytometric analysis of PAN RNA expression in multiple conditions, from cells incubated with no probe (left), or cells subjected to hybridization using a probe for PAN RNA, comparing virus-negative BL41 cells (second from left) with KSHV+ BCBL-1 cells that were either untreated or stimulated (with TPA and sodium butyrate (NaB)) for 72 hours. Data depict lymphocytes that were singlets, defined by sequential removal of doublets. Representative images were defined as samples that were closest to the median frequency. Quantification of (B) the frequency of PAN RNA+ cells and (C) PAN RNA median fluorescence within PAN RNA+ cells, comparing untreated or stimulated BCBL-1 cells. D, E) Flow cytometric analysis of untreated BCBL-1 cells, using histogram overlays, to compare cell size (forward scatter, FSC), granularity (side scatter, SSC) and ORF73 expression in cells that were either PAN RNA negative [-] (blue line) or PAN RNA positive [+] (red line). Data show (D) histogram overlays of these populations with fluorescence quantification provided in panel E. ORF73 analysis includes samples in which there was no ORF73 probe (i.e. “No Probe”, in solid gray), to define background fluorescence. (F, G) Flow cytometric analysis of stimulated BCBL-1 cells, using histogram overlays, to compare cell size (forward scatter, FSC), granularity (side scatter, SSC) and ORF73 expression in cells that are either PAN RNA negative [-] (blue) or PAN RNA positive [+] (red). Data show (F) histogram overlays of these populations with fluorescence quantification provided in panel G, with ORF73 analysis including a “No Probe” sample (gray) to define background fluorescence. Due to variable baseline fluorescence values for SSC and ORF73 between experiments, values were internally standardized to fluorescent intensities within the PAN RNA negative population for each experiment, with data depicting mean ± SEM. Symbols in panels B and C indicate values from individual samples. Data are from two independent experiments, with biological replicates within each experiment, with total number of biological replicates as follows: No Probe (n = 2), BL41 control (n = 3), BCBL-1 untreated (n = 6), BCBL-1 stimulated (n = 6). Statistical analysis was done using an unpaired t test with statistically significant differences as indicated, * p<0.05, ** p<0.01, **** p<0.0001.

We next analyzed the properties of BCBL-1 cells as a function of PAN RNA expression. In untreated cells, PAN RNA+ or RNA- cells had comparable cell size (define by forward scatter, FSC) and granularity (defined by side scatter, SSC). ORF73 RNA expression was low in untreated BCBL-1 samples, with signal intensity in PAN RNA- cells close to the background fluorescence observed in unstained samples. PAN RNA+ cells in untreated cultures had a modest increase in ORF73 RNA expression relative to PAN RNA- cells (Fig. 3D-E). In stimulated BCBL-1 cultures, PAN RNA+ cells had a modest decrease in cell size (defined by forward scatter) and a trend towards reduced granularity (defined by side scatter) compared to PAN RNA- cells (Fig. 3F-G). Stimulated BCBL-1 cultures also had an increased ORF73 signal when compared to unstained samples (Fig. 3F), with PAN RNA+ cells again showing ~2-fold increase compared to PAN RNA- cells (Fig. 3F-G). These data demonstrate robust detection of PAN RNA by PrimeFlow™, and further identify PAN RNA expression in a subset of both untreated and reactivation-induced BCBL-1 cells.

### Detection of endogenous viral gene expression during EBV latency and reactivation

EBV encodes two abundant non-coding RNAs, the EBV-encoded RNAs (EBERs) EBER1 and EBER2. We tested the ability of the PrimeFlow™ method to detect EBER in an EBV positive, Burkitt lymphoma type I latency cell line, Mutu I [23]. EBER expression was detected in ~45% of Mutu I cells in either untreated or TPA stimulated conditions, with EBER+ cells defined relative to background probe hybridization in the KSHV- and EBV-negative BL41 cell line, a conservative measurement (Fig. 4A). TPA stimulated Mutu I cells showed a modest, 2-fold increase in EBER expression on an individual cell basis, relative to untreated EBER+ cells (Fig. 4B). Based on these data, EBER expression in Mutu I cells appears to be constitutive, with stimulation under these conditions resulting in minimal consequences on either the frequency or per-cell expression of the EBERs.

**Figure 4.**
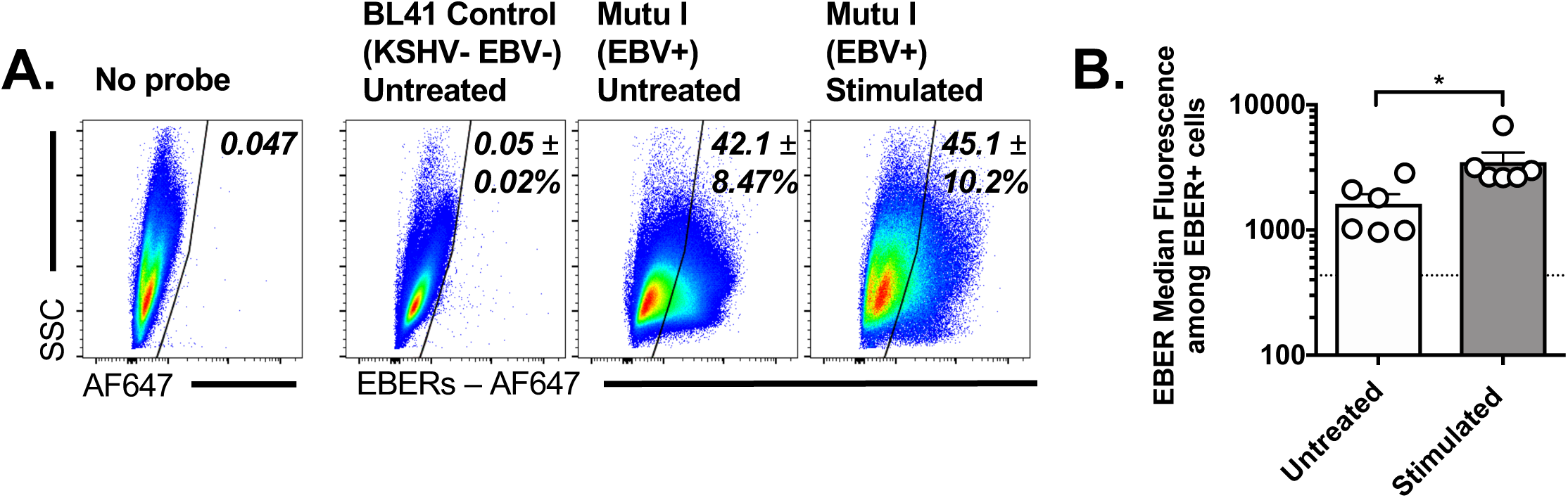
Single-cell analysis of EBV EBER expression in the Mutu I B cell lymphoma cell line. (A) Flow cytometric analysis using the PrimeFlow™ method, to quantify EBER expression in multiple conditions, from cells incubated with no probe (left), or cells subjected to hybridization using a probe for the EBERs, comparing virus-negative BL41 cells (second from left) with Mutu I EBV+ cells that were either untreated (with DMSO) or stimulated (with TPA in DMSO) for 48 hours. Data depict lymphocytes that were singlets, defined by sequential removal of doublets. Representative images were defined as samples that were closest to the median frequency, with data depicting mean ± SEM. (B) Quantitation of median EBER fluorescence within EBER+ Mutu I cells in untreated and stimulated cultures. Horizontal dashed line indicates the background fluorescent signal from BL41 controls. Data are from two independent experiments, with biological replicates within each experiment, with total biological replicates as follows: No Probe (n = 2), BL41 Control untreated (n = 6), Mutu I untreated (n = 6), Mutu I stimulated (n = 6). Statistical analysis was done using an unpaired t test with statistically significant differences as indicated, * p<0.05.

To extend these findings, we further analyzed EBER expression in a panel of LCLs and during in vitro infection of primary human B cells. EBER expression was readily detected in LCL cultures, with EBER+ cells also found in a subset of human primary B cells following in vitro EBV infection (Supplemental Fig. 2, 3). During EBV infection of human primary B cells, EBER+ B cells had an increased cell size and granularity relative to EBER- B cells, with increased expression of both CD69 and actin mRNA, an activated cell phenotype (Supplemental Fig. 3D). These data demonstrate cell to cell variation in EBER expression and suggest EBER expression as a potential discriminator to investigate variability during EBV infection.

### Single-cell analysis of actin mRNA degradation as a readout of virus-induced host shutoff

Many herpesviruses, including γHV68, EBV and KSHV, induce host shutoff during lytic replication and reactivation from latency, a process characterized by dramatic decreases in host mRNAs [24, 25]. Consistent with published reports [24], qRT-PCR analysis of a cellular housekeeping gene, β-actin (Actb), showed reduced actin mRNA in γHV68 lytically-infected fibroblasts by 18 hours pi (Fig. 5A). While mock-infected samples had a uniformly positive population of actin RNA^high^ cells detectable by PrimeFlow™, γHV68-infected fibroblast cultures demonstrated a bimodal distribution of actin RNA^high^ and actin RNA^low^ cells (Fig. 5B). The actin RNA^low^ population had a fluorescent signal that was only modestly above background fluorescence (defined by the “No probe” sample), suggesting an all-or-none phenomenon in which cells either had no change in actin RNA levels or had pronounced actin RNA degradation. Simultaneous analysis of TMER and actin RNA expression revealed that actin RNA^low^ cells were frequently TMER^high^, with actin^high^ cells frequently TMER^negative^ at this time (Fig. 5C).

**Figure 5.**
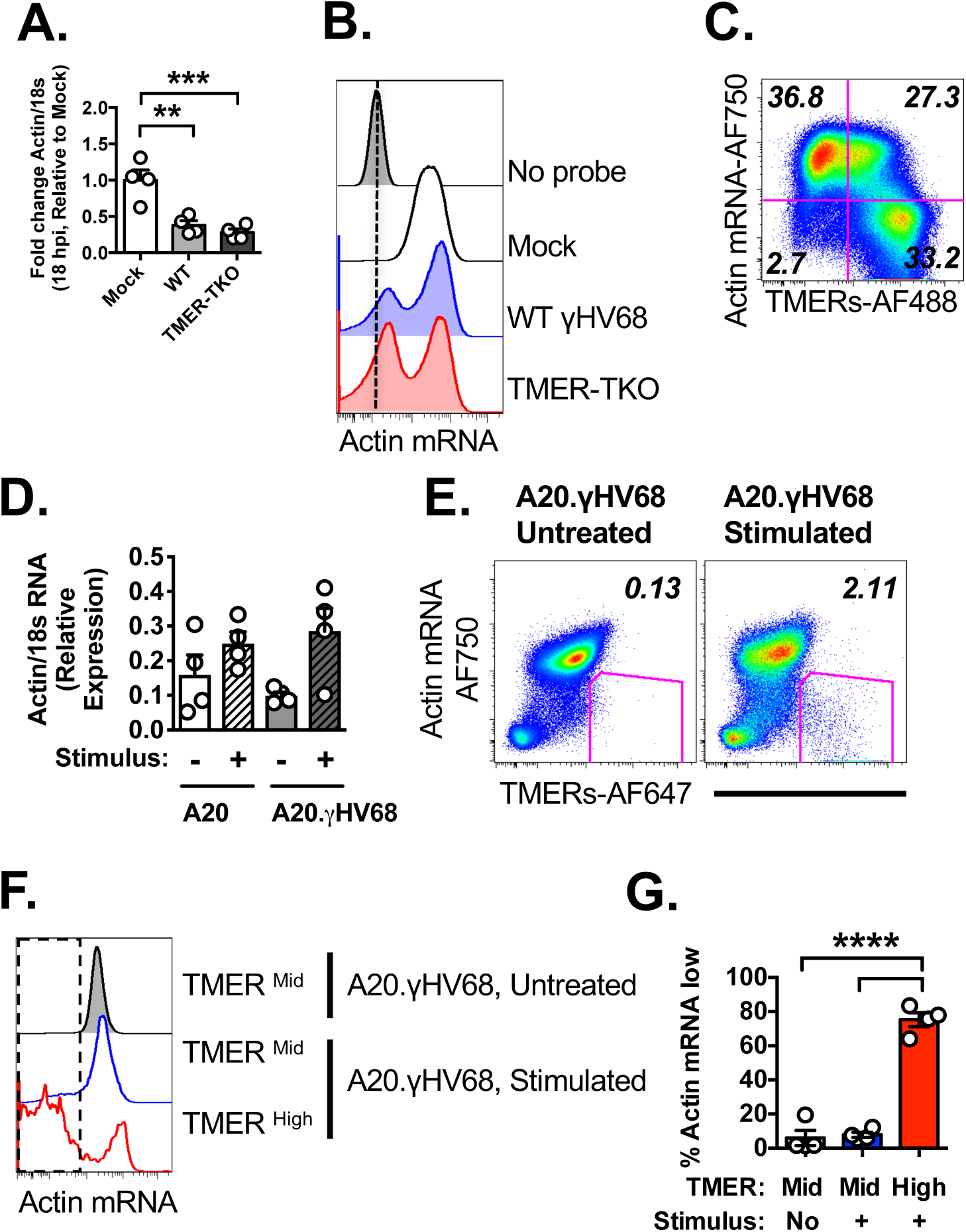
Actin mRNA degradation identifies virally-infected cells experiencing virus-induced host shutoff. Actin mRNA analysis by qRT-PCR (A,D) or by flow cytometric analysis using PrimeFlow™ (B,C,E-G), comparing cells with variable infection status. (A) qRT-PCR analysis of beta-actin (Actb) mRNA expression relative to 18s RNA in mock, WT or TMER-TKO infected 3T12 fibroblasts at 18 hpi. (B) PrimeFlow™ analysis of actin mRNA in 3T12 fibroblasts, either unstained (“No probe”), mock-infected or infected with WT γHV68 or TMER-TKO at 18 hpi. (C) PrimeFlow™ analysis of actin mRNA and TMER expression in WT γHV68 infected fibroblasts at 18 hpi. (D) qRT-PCR analysis of beta-actin (Actb) mRNA expression relative to 18s RNA in A20, virus-negative cells and A20.γHV68 (HE2.1) cells, untreated or stimulated with TPA for 24 hrs. (E) PrimeFlow™ analysis of actin mRNA and TMER expression in untreated and stimulated A20.γHV68 cells, with the frequency of TMER^high^ actin mRNA^low^ cells indicated, based on the gated events. (F,G) Actin mRNA analysis by PrimeFlow™ using either (F) histogram overlays or (G) quantifying frequencies, comparing A20.γHV68 cells that were either untreated or stimulated, further stratified by whether the cells were TMER^mid^ or TMER^high^ (using the gating strategy defined in Fig. 2C). All flow cytometry data depict single cells, defined by sequential removal of doublets. Data are from two-three independent experiments, with biological replicates within each experiment. Graphs depict the mean ± SEM, with each symbol identifying data from a single replicate. Statistical analysis was done using one-way ANOVA, subjected to Tukey’s multiple comparison test (A, D, G), with statistically significant differences as indicated, ** p<0.01, *** p<0.001, **** p<0.0001.

To determine whether actin RNA regulation could also be observed during γHV68 latency and reactivation, we measured actin RNA levels in A20.γHV68 cells. Parental, virus-negative A20 cells and A20.γHV68 cells had relatively comparable actin RNA levels by qRT-PCR, in both untreated and stimulated cells (Fig. 5D). Given that host shutoff is expected to primarily occur in rare, reactivating cells, we measured actin RNA degradation relative to TMER expression by the PrimeFlow™ method. Untreated A20.γHV68 cultures had no discernable population of TMER+ actin RNA^low^ events, whereas stimulated cultures were characterized by a rare population of TMER^high^ actin RNA^low^ cells (Fig. 5E). We further compared actin RNA expression between TMER^mid^ and TMER^high^ cells, in untreated versus stimulated cultures using our previously defined subpopulations (Fig. 2). While TMER^mid^ cells from either untreated or stimulated cultures were predominantly actin RNA+, TMER^high^ cells from stimulated cultures showed a significant increased frequency of actin RNA^low^ events (Fig. 5F-G). These studies reveal actin RNA as a sensitive indicator of virus-induced host shutoff, and demonstrate this as an all-or-none phenomenon that can be readily queried at the single-cell level.

### Heterogeneity of gene expression during de novo lytic replication

Next, we revisited our analysis of gene expression during de novo lytic infection of fibroblasts, to examine co-expression relationships between viral ncRNA (TMERs), viral mRNA (the γHV68 ORF73), viral protein (RCA protein) and cellular actin mRNA degradation [19, 24]. Mouse 3T12 fibroblasts were infected with a multiplicity of infection (MOI=5 plaque forming units of virus/cell), harvested 16 hpi and then subjected to the PrimeFlow™ method.

To enable an unbiased, automated analysis of gene expression profiles in γHV68 lytically infected cells relative to mock infected cells, data were subjected to the automated clustering algorithm X-shift [26], to identify potential subpopulations of cells with heterogeneous gene expression in these cultures. By sampling 1,000 cells from multiple mock- and virus-infected cultures, the X-shift algorithm consistently identified 7 major clusters of cells (Fig. 6A) defined by varying gene expression patterns. While some of the clusters were exclusively found in mock-infected cultures, virus-infected cultures contained three broad types of cell clusters: 1) cells, with no detectable expression of either the TMERs or ORF73 and normal actin RNA, 2) fully infected cells, with robust expression of the TMERs, ORF73, actin RNA downregulation and frequent expression of the RCA protein, and 3) intermediate populations characterized by variable expression of the TMERs and ORF73 (Fig. 6A).

**Figure 6.**
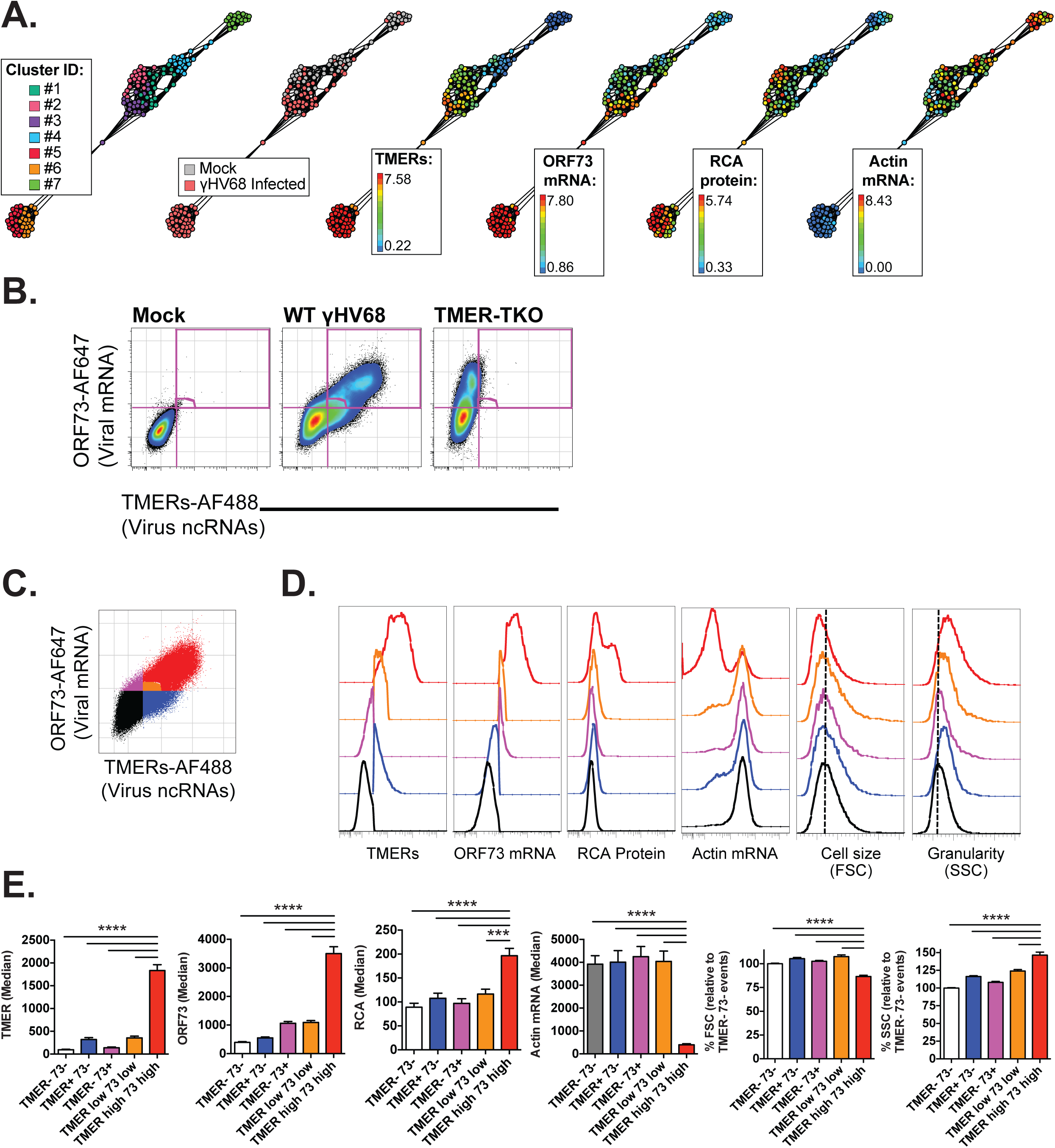
Heterogeneous viral gene expression at the single-cell level during lytic replication. Viral, host RNA flow cytometric analysis in γHV68-infected fibroblasts at 16 hpi defined by the PrimeFlow™ method, comparing (A) X-shift clustering analysis and (B-E) biaxial gating analysis for the indicated features. (A) Automated, clustering analysis using the X-Shift algorithm on 10,000 events total, compiled from mock- and γHV68-infected fibroblasts at 16 hrs pi (1,000 events randomly imported per sample, mock infected n=4, γHV68-infected n=6) identifies multiple clusters of cells with differential gene expression (7 clusters, colored distinctly, “Cluster ID”), with these clusters then depicted for expression of TMERs, ORF73, Actin mRNA, and RCA. Range of expression is identified for each parameter. (B) Analysis of TMER and ORF73 co-expression in mock (left), WT γHV68-infected (middle), and TMER-TKO-infected (right) samples, with gates depicting populations with different gene expression profiles, defined relative to mock and TMER-TKO infected samples. (C) Color-coded populations from WT-infected sample in panel B, with each color indicating a different gene expression profile. (D) Histogram overlays of the five populations identified in panel C for the indicated parameters. (E) Quantitation of gene expression among the five populations identified in panel C, using the same color-coding strategy. Data are from three independent experiments, with each experiment containing biological replicates. Flow cytometry data shows single cells that are DNA+ (DAPI+). Statistical significance tested by one-way ANOVA, comparing the mean of TMER^high^ ORF73^high^ cells to all other means, followed by Dunnett’s multiple testing correction. Significance identified as *** p<0.001, **** p<0.0001.

To validate these findings using a more conventional method, we compared TMER and ORF73 RNA co-expression on a biaxial plot. By comparing mock-infected, WT-infected and TMER-TKO-infected cultures, this analysis revealed five populations of gene expression (Fig. 6B), including cells with: 1) no detectable expression of either viral RNA (TMER- ORF73-), 2) TMER+ ORF73- cells (bottom right quadrant), 3) TMER- ORF73+ cells (upper left quadrant), 4) TMER^low^ ORF73^low^ cells (lower left edge of the upper right quadrant), and 5) TMER^high^ ORF73^high^ cells (upper right quadrant). The definition of TMER positive events was defined based on background fluorescent levels observed in TMER-TKO infected cultures (Fig. 6B). These 5 populations were each assigned a unique color for subsequent analysis (Fig. 6C).

We compared cellular phenotype and gene expression across these 5 populations. Analysis of TMERs, ORF73, actin RNA, RCA protein, cell size (forward scatter), and granularity (side scatter) revealed multiple types of viral gene expression. TMER- ORF73- cells (in black) had no evidence of viral gene expression, with no detectable viral protein (RCA) or actin downregulation (Fig. 6D-E). Cells with low expression of either the TMERs and/or ORF73 contained viral RNAs, but had minimal expression of either viral protein or actin downregulation (Fig. 6D-E). In stark contrast, cells that were TMER^high^ ORF73^high^ (in red, Fig. 6D-E) had multiple characteristics of progressive virus infection including a prominent fraction of cells that expressed RCA and/or had actin RNA downregulation. Further, TMER^high^ ORF73^high^ cells were consistently smaller in cell size (defined by forward scatter, FSC) and higher in granularity (defined by side scatter, SSC), a feature that was unique to this phenotype (Fig. 6D-E).

TMER^high^ ORF73^high^ also demonstrated robust expression of an early gene (ORF64) and a late gene (ORF18) (Supplemental Fig. 4A). While some TMER^low^ ORF73^low^ cells expressed early and late genes, cells defined as TMER- 73- had no discernable expression of either early or late genes (Supplemental Fig. 4A). These data emphasize that a subset of 3T12 fibroblasts subjected to infection with an MOI=5 and harvested at 16 hpi have negligible signs of viral gene expression using this method. This is further illustrated by comparing mock and γHV68 infected cultures for TMER, ORF73, ORF64 and ORF18 expression which demonstrates a subset of cells with minimal expression across each of these viral genes (Supplemental Fig. 4B). One potential explanation for this could be that TMER- ORF73- cells are not infected. To test this, we sort purified cells based on TMER and ORF73 expression, followed by quantitation of viral DNA. TMER^high^ ORF73^high^ cells and TMER+ 73- cells had comparable levels of viral DNA, which were approximately ten-fold higher than viral DNA levels in TMER- 73- cells (Supplemental Fig. 4C). Despite this moderate difference in viral DNA, however, viral DNA levels in TMER- 73- cells were still significantly above background signal present in mock infected cultures (Supplemental Fig. 4C). Another potential explanation for this limited gene expression in TMER- 73- cells is that perhaps they are in a distinct phase of the cell cycle relative to other cells. We found that there was no sizable impact of cell cycle stage on whether cells were TMER- ORF73- or TMER^high^ ORF73^high^ (Supplemental Fig. 4D). These data identify unexpected heterogeneity during in vitro lytic infection, with TMER^high^ ORF73^high^ cells characterized by robust viral transcription, TMER^low^ ORF73^low^ cells characterized by a lower frequency of cells expressing early and late genes, and TMER- ORF73- cells containing viral DNA with little to no detectable viral gene expression.

Given the heterogeneous patterns of RNA and protein expression among lytically-infected cells, we next queried TMER subcellular localization as a function of viral gene expression using imaging flow cytometry. While the majority of TMER+ cells had a primarily nuclear TMER localization (defined by DAPI co-localization, as in [27]), the frequency of cells with nuclear TMER localization was highest among TMER+ ORF73- cells and lowest among TMER+ ORF73+ RCA+ cells (Fig. 7A-B). These data suggest that the TMERs can be localized in either the nucleus or cytoplasm during γHV68 lytic replication, and that this localization is not strictly a function of magnitude of gene expression.

**Figure 7.**
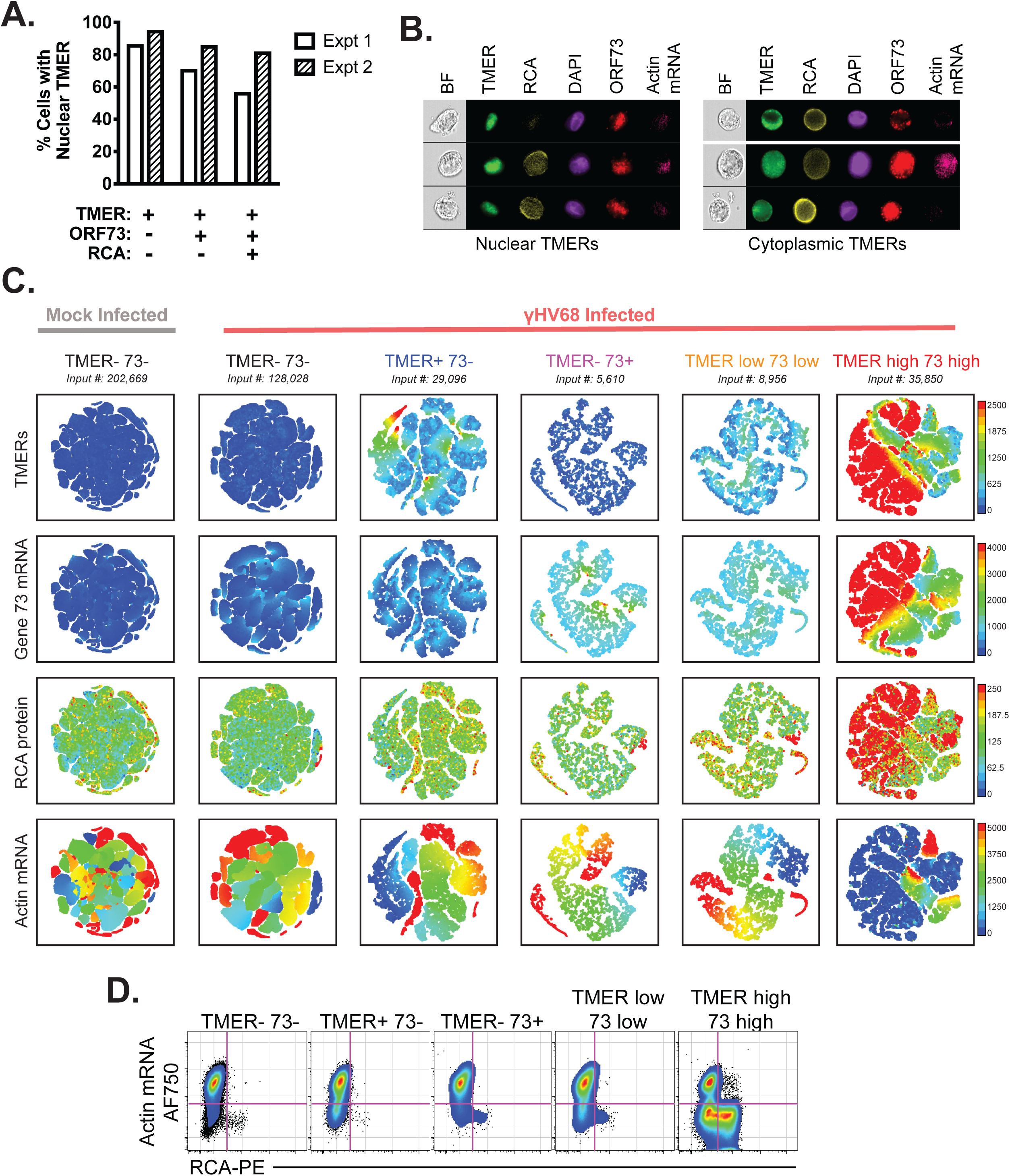
Lytic replication is characterized by heterogeneous TMER localization and variable penetrance of actin RNA degradation. Viral, host RNA flow cytometric analysis in γHV68-infected fibroblasts at 16 hpi defined by the PrimeFlow™ method. (A) The frequency of γHV68-infected fibroblasts with TMERs primarily in the nucleus was quantified by ImageStream, with data showing the frequency of cells in which TMER:DAPI colocalization (i.e. similarity score) was >1. (B) Images showing brightfield (BF), TMER, RCA protein (RCA), DAPI, ORF73 and actin mRNA localization, comparing cells with nuclear TMER localization (left) versus cytoplasmic TMER localization (right). (C) Analysis of cell subpopulations stratified by TMER and ORF73 expression (defined in Fig. 6C), subjected to the tSNE dimensionality reduction algorithm. Data and cell populations are derived from the dataset presented in Fig. 6, showing all DNA+ (DAPI+) single cells (FSC-A, SSC-A) subjected to the tSNE algorithm. The tSNE algorithm provides each cell with a unique coordinate according to its expression of Actin mRNA, RCA, ORF73, and TMERs, displayed on a two-dimensional plot (tSNE1 versus tSNE2). Visualization grid of tSNE plots, with plots arranged according to marker expression (rows) relative to phenotype of the cellular population examined (columns). (D) Biaxial analysis of actin RNA versus RCA protein, among the five populations identified in Fig. 6. Flow cytometry data shows single cells that are DNA+ (DAPI+). Data in panels A-B from two independent experiments, in panel C-D from three independent experiments.

Finally, we used tSNE, a dimensionality reduction algorithm, to better delineate the relationship between TMER, ORF73, RCA protein and actin downregulation across populations defined by variable TMER and ORF73 expression. In tSNE-based visualization, events are subjected to dimensionality reduction, with all events plotted according to the composite parameters tSNE1 and tSNE2. In each tSNE-based plot, values from individual cells are depicted as individual dots on the plot. Cells with similar expression profiles are visually clustered together using this algorithm (e.g. [28]). Consistent with our histogram analysis (Fig. 6D-E), cells that were negative for TMER and ORF73 and cells that expressed either TMERs or ORF73 were relatively uniform in gene expression (Fig. 7C). In contrast, TMER^high^ ORF73^high^ cells expressed a wider array of phenotypes, including both a predominant fraction of cells that were actin RNA^low^ RCA+, and a distinct group of cells that were actin RNA+ RCA- (Fig. 7C). Notably, RCA expression and actin degradation were inversely correlated, with very few cells that expressed RCA also high for actin RNA. Actin RNA+ populations among TMER^high^ ORF73^high^ cells were associated with larger cell size (Supplemental Fig. 5). The diversity of phenotypes among TMER^high^ ORF73^high^ cells was confirmed by biaxial gating of actin RNA versus RCA protein expression (Fig. 7D). In total, these data indicate heterogeneous progression of lytic replication in vitro. While some cells have robust viral mRNA and protein expression, additional cell subsets are characterized by limited or divergent gene expression.

### Lytic cycle viral gene expression is influenced by target cell type, changes over time, and is modulated by conditions that alter lytic replication

Our previous studies of γHV68 lytic replication used 3T12 fibroblasts, an immortalized cell line. Mouse embryonic fibroblasts (MEFs) are primary cells that are highly permissive for γHV68 infection, with 5 to 10-fold greater sensitivity to virus infection than 3T12 fibroblasts [29]. We therefore compared viral gene expression between 3T12 fibroblasts and MEFs, using MEFs derived either from wild-type or IFNAR1 KO mice. All cells were infected with WT γHV68 (MOI=5) and analyzed at 16 hpi. Both WT and IFNAR KO MEFs showed enhanced viral gene expression relative to infected 3T12 cells at 16 hpi, with infected MEF cultures characterized by an increased frequency of events that expressed ORF73, TMERs, RCA and were actin^low^ (Fig. 8A). Infected MEFs also had an increased frequency of ORF73+TMER+ cells and a decreased frequency of ORF73-TMER- cells relative to 3T12 cells (Fig. 8B). When we analyzed the cellular distribution of viral gene expression across all 16 possible combinations of viral gene expression, resulting from different combinations of ORF73, TMER, RCA and actin^low^ phenotypes, infected MEF cultures compared to infected 3T12 cultures had: i) fewer ORF73-TMER- cells, ii) an increased frequency of ORF73+TMER+actin^low^ cells, iii) with the most frequent subset defined as cells with full viral gene expression (i.e. ORF73+TMER+RCA+actin^low^) (Fig. 8C). Viral gene expression distribution between WT and IFNAR1 KO MEFs were generally comparable, suggesting little to no contribution of type I IFN receptor mediated signaling as a regulator of viral gene expression in MEFs. These data demonstrate that the diversity and relative abundance of viral gene expression can be influenced by target cell type.

**Figure 8.**
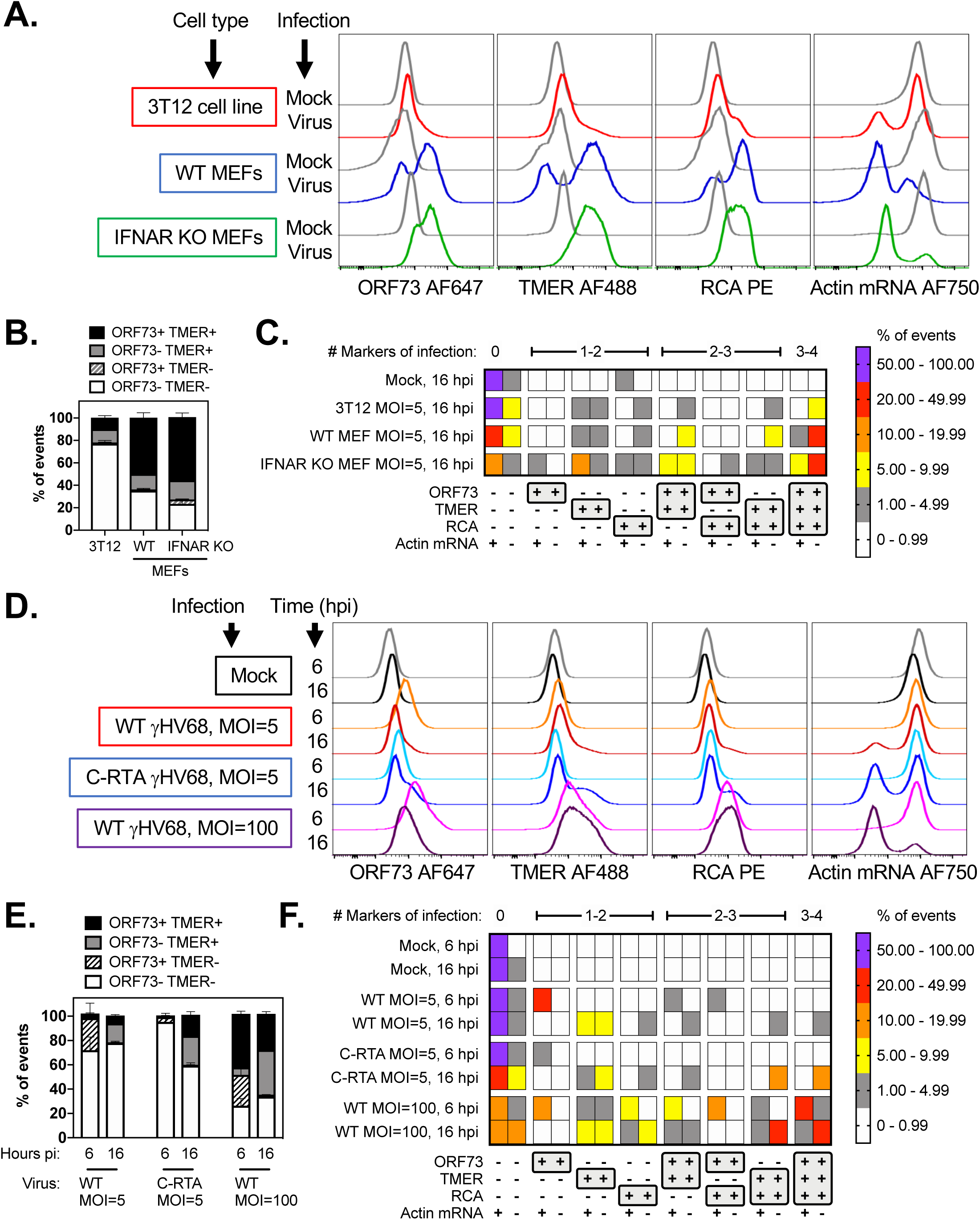
Lytic viral gene expression is influenced by target cell type, changes over time, is enhanced by conditions which promote lytic replication. Fibroblasts (3T12 cells or mouse embryonic fibroblasts, MEFs) were infected with WT or C-RTA γHV68 at 5 or 100 PFU/cell, and harvested at either 6 or 16 hpi as indicated. (A-C) Analysis of viral gene expression by PrimeFlow™ in either 3T12 or MEFs following WT γHV68 infection (MOI=5) and harvested at 16 hpi. (A) Analysis of viral gene expression by PrimeFlow™, comparing gene expression by histogram overlays between 3T12, WT MEFs and IFNAR KO MEFs. (B) The frequency of events stratified based on expression of ORF73 and TMER, or (C) based on all 16 possible combinations of gene expression with the frequency of events in each population plotted as a heatmap. Data are from three independent experiments harvested at the indicated time, subjected to fluorescent barcoding, pooled together and subjected to PrimeFlow™ analysis in a single analysis. (D-F) 3T12 fibroblasts were infected with WT or C-RTA γHV68 at 5 or 100 PFU/cell, and harvested at either 6 or 16 hpi as indicated. (D) Analysis of viral gene expression by PrimeFlow™, comparing gene expression by histogram overlays between different virus infection conditions and times post-infection, as indicated. (E) The frequency of events stratified based on expression of ORF73 and TMER, or (F) based on all 16 possible combinations of gene expression with the frequency of events in each population plotted as a heatmap. Data are from three independent experiments harvested at the indicated time, subjected to fluorescent barcoding, pooled together and subjected to PrimeFlow™ analysis in a single analysis. Data depict mean values, with SEM plotted in panels B, D, and heatmap values colored based on the categorical key provided in the figure. Flow cytometry data depict single cells, defined by sequential removal of doublets according to SSC-A x SSC-H and FSC-A x FSC-H, after which files were subjected to debarcoding to identify each of the independent samples in the analysis.

The increased viral gene expression observed in MEFs raised the possibility that viral gene expression could be further modulated in 3T12 cells, either as a function of time, multiplicity of infection or viral genotype. We therefore compared viral gene expression in 3T12 fibroblasts, comparing: i) WT γHV68 infection with 5 PFU/cell, ii) C-RTA γHV68 (a recombinant virus engineered to overexpress Rta, the immediate early viral transactivator [30]) infection with 5 PFU/cell, and iii) WT infection using 100 PFU/cell. Cells were harvested at 6 and 16 hpi. When comparing WT infection at MOI=5 between 6 and 16 hpi, ORF73 expression peaked at 6 hpi with decreased expression by 16 hpi, in contrast to TMER, RCA and actin^low^ phenotypes which were infrequent at 6 hpi and increased by 16 hpi (Fig. 8D). 3T12 cells subjected to WT infection at an MOI=100 showed an increased expression of ORF73, TMER and RCA by 6 hpi, above WT infection with an MOI=5. By 16 hpi, cultures infected with C-RTA and WT MOI=100 had increased frequencies of TMER+ and ORF73+TMER+ events relative to WT MOI=5 cultures (Fig. 8E).

The diversity and progression of viral gene expression was further illustrated by analyzing the frequency of all 16 different combinations of possible gene expression between these different virus conditions. While WT MOI=5 cultures showed a time-dependent switch from ORF73+ events to TMER+, ORF73+TMER+, and ORF73+TMER+RCA+actin^low^ events, the frequency of each of these populations was typically below 10% of cultures (Fig. 8F). In contrast, cultures infected with C-RTA had limited viral gene expression at 6 hpi, but showed enhanced progression to both TMER+RCA+actin^low^ and ORF73+TMER+RCA+actin^low^ phenotypes relative to WT infection MOI=5 (Fig. 8F). Cultures infected with WT virus at an MOI=100 showed rapid induction of multiple phenotypes, including a sizable portion of cultures which were ORF73+RCA+ and ORF73+TMER+RCA+ by 6 hpi, both populations which retained actin RNA expression. By 16 hpi, cultures infected with WT virus at an MOI=100 had sizable frequencies of TMER+RCA+actin^low^ and ORF73+TMER+RCA+actin^low^ events (Fig. 8F). While C-RTA and MOI=100 infected cultures showed enhanced viral gene expression, WT cultures treated with phosphonoacetic acid (PAA), an inhibitor of viral DNA synthesis, showed impaired expression of TMER with a complete absence of TMER^high^ events, negligible ORF73 expression and a pronounced inability to induce an actin^low^ phenotype (Supplemental Fig. 6). These data demonstrate diversity of viral gene expression that can occur during lytic cycle and provide direct demonstration that these different gene expression profiles can be experimentally manipulated by either enhancing or restraining lytic replication.

## DISCUSSION

Herpesvirus gene expression has been historically analyzed in bulk cell populations. These studies have provided an essential cornerstone to understanding the transcriptional and translational capacity of the herpesviruses. Despite this, recent studies on cellular and viral transcription from other systems have emphasized a high degree of cell-to-cell variation in gene expression [6-11, 13, 14], something we have further investigated here. By applying the PrimeFlow™ methodology to measure endogenous viral gene expression across multiple gammaherpesviruses, and multiple stages of infection, we have gained critical new insights into the inter-relationships of gene expression at the single-cell level.

A primary focus of the current study has been to analyze expression of γHV ncRNAs. Although the TMERs, EBERs and PAN RNA all represent abundant γHV ncRNAs, these ncRNAs are transcribed by distinct mechanisms: KSHV PAN is a highly-inducible, RNA pol II-transcribed ncRNA [20], in contrast to the RNA pol III-transcribed TMERs and EBERs [15, 31]. This differential regulation was mirrored in the expression patterns we observed. Whereas TMERs and EBERs were detected in a large fraction of latently infected cells, PAN RNA was expressed in a low frequency of latently infected cells, with prominent induction following cell stimulation and the induction of reactivation. The viral ncRNAs were efficiently detected, as might be predicted due to their abundance. The viral ORF73 encodes a transcription factor that is expressed at a far lower level and are also efficiently detected, demonstrating that rare mRNAs can be measured coincidently with abundant RNAs and with proteins, with no modifications required. A unique advantage of our current approach is the ability to measure the frequency of ncRNA expressing cells and changes in expression within individual cells. This has been particularly insightful for the identification of rare PAN RNA+ cells in untreated BCBL-1 cells and a TMER^high^ subpopulation of cells in reactivating A20.γHV68 cells. Integrating this method with cell sorting will afford future opportunities to investigate unique properties of these rare cell populations.

Among the viral ncRNAs measured, in-depth analysis of TMER expression during γHV68 infection has revealed new insights into infection. In the context of latency, the TMERs are constitutively expressed in many, but not all, latently infected cells using the A20.γHV68 model. Further, stimulating these cells to undergo reactivation has a minimal effect on the frequency of cells expressing intermediate levels of TMERs (i.e. TMER^mid^ cells), instead resulting in the appearance of a minor population of TMER^high^ cells. Notably, TMER^high^ cells show additional features of lytic cycle progression, including actin RNA degradation and RCA protein expression. Why only some latently infected cells show the TMER^high^ phenotype, and what regulates the inducible expression of the RNA pol III-transcribed TMERs remain important questions raised by this analysis.

Of the γHVs studied here, only γHV68 has a robust in vitro lytic replication system. Our studies on γHV68 lytic replication revealed multiple unanticipated results. First, our analysis identified heterogeneity of viral gene expression, stratified by differential viral gene expression of the TMERs and ORF73. Strikingly, in cultures subjected to infection with 5 PFU per cell, conditions generally considered to induce synchronous lytic replication, there were many cells with limited viral gene expression, expressing low levels of either the TMERs and/or ORF73, but lacking additional signs of virus gene expression (i.e. actin RNA degradation or RCA protein expression). This analysis further identified a reproducible subset of cells with negligible viral gene expression, defined by a TMER- ORF73- phenotype. Curiously, these cells contain viral DNA but did not have further evidence of viral gene expression. Whether these cells represent an abortive state of infection, or if viral genes are expressed at levels below our current limit of detection remains to be determined. Our studies further revealed that only some viral RNA+ cells showed full progression of virus infection characterized by robust viral gene expression, defined as TMER^high^ ORF73^high^ RCA+ actin RNA^low^. Though progression to full viral gene expression was enhanced by infection of primary fibroblasts, infection with the C-RTA virus, or a significantly higher MOI, even in these conditions there remained heterogeneity in viral gene expression. Whether heterogeneity of viral gene expression persists when cells manifest cytopathic effect, a process which occurs after 16 hpi, is not yet known. While there is precedence that reactivation from latency in KSHV infection can be asynchronous [9], this heterogeneity of viral gene expression during in vitro lytic replication was unanticipated and suggests that lytic infection under these reductionist conditions is either asynchronous, abortive, or inefficient. This heterogeneity of gene expression raises important questions regarding the universality of the prototypical cascade of immediate early, early and late gene expression that is widely accepted in the herpesvirus field and suggests additional levels of complexity that may be obscured by bulk cell analysis. The molecular mechanisms that are responsible for this heterogeneity still remain to be elucidated but appear to be independent of cell cycle stage with little to no contribution of type I interferon receptor mediated signaling based on relatively comparable gene expression profiles in wild-type and IFNAR1 KO MEFs.

This method allows multiplexed analysis of single-cell gene expression, to both directly measure viral RNAs and downstream consequences of gene expression including viral protein production and host RNA degradation, secondary to protein translation. This approach has notable advantages to conventional analyses of gene expression: 1) it can measure endogenous viral gene expression (both mRNA and ncRNA) in the absence of recombinant viruses or marker genes, and 2) it can rapidly analyze gene and protein expression inter-relationships, across millions of cells, providing unique complementary strengths to other single-cell methodologies (e.g. single-cell RNA-seq). In the future, this method can be further integrated with additional antibody-based reagents, to simultaneously query post-translational modifications (e.g. protein phosphorylation) as a function of cell cycle stage. It is also notable that through the use of imaging flow cytometry, it is possible to interrogate subcellular RNA and protein localization throughout distinct stages of virus infection, studies revealing single cell variability in subcellular RNA localization. We anticipate that this approach will have widespread utility, from addressing fundamental questions about herpesvirus gene expression using reductionist approaches, to a better delineation of replication or reactivation defects in viral mutants, to the in vivo identification of viral gene expression in primary infected samples.

The approach presented here provides a powerful complementary method to other single cell methods, affording the opportunity to query a diverse set of experimental manipulations in a relatively rapid manner. It is worthwhile to note, however, some important considerations with this approach. First, these studies rely on fluorescent measurements of probe hybridization using a flow cytometer. Though this approach detects a range of viral and host RNAs, sensitivity of this method is influenced by target RNA expression level, fluorophores used for the analysis and controls to define probe specificity and limit of detection. Ideally, comparisons can be strengthened by comparing isogenic conditions (e.g. comparing WT virus with a genetically deficient virus, as done with the TMER deficient virus). In cases where non-isogenic conditions are used (e.g. comparing across cell lines), variable background fluorescence may limit the sensitivity of this method. For new users of this technology, we strongly suggest the use of multiple controls, including a full-minus-one (i.e. FMO) control to accurately define background and signal to noise ratio for targets of interest. We consider multiparameter flow cytometry, as well as mass cytometry [32, 33], two increasingly useful technologies to afford new insight into heterogeneity of virus infection and gene expression that provide a complementary approach to other single cell technologies.

In total, these studies demonstrate the power of single-cell analysis of herpesvirus gene expression. Our data emphasize the heterogeneity of γHV gene expression at the single-cell level, even in conditions considered to result in uniform infection. The factors that underlie this heterogeneity are currently unknown, but could reflect either asynchronous or inefficient infection in many infected cells (e.g. in the context of lytic infection). The existence of specific infected cell subsets, based on heterogeneous gene expression, may identify new susceptibilities and points for intervention during the course of virus infection. Whether this variation arises from viral or cellular heterogeneity is a fundamental question for future research.

## MATERIALS AND METHODS

### Viruses and tissue culture

γHV68 viruses were derived from the γHV68 strain WUMS (ATCC VR-1465) [34], using either bacterial artificial chromosome-derived wild-type (WT) γHV68 or γHV68.TMER-Total KnockOut (TMER-TKO) [16], or virus derived by homologous recombination, γHV68.C-RTA [30]. Virus stocks were passaged, grown, and titered as previously described [16]. Mouse 3T12 fibroblasts (ATCC CCL-164) were infected with a multiplicity of infection (MOI) of either 5 or 100 plaque forming units/cell, analyzed 6-18 hpi. Primary mouse embryonic fibroblasts isolated from C57BL/6 mice (B6 MEFs) or from B6 IFNAR1 KO mice (IFNAR KO MEFs, kindly provided by Dr. Thomas E. Morrison at the University of Colorado) were cultured in DMEM with 10% FBS, 1% Penicillin/Streptomycin and then either mock- or virus-infected (MOI=5) harvested at 16 hpi. For experiments using PAA, PAA (Sigma-Aldrich, catalog #284270) was added at a concentration of 200 μg/mL at time of virus infection and left on cultures until the time of harvest. The parental, virus-negative A20 B cell lymphoma cell line was obtained from ATCC (ATCC^®^ TIB-208^™^) and cultured in RPMI 1640 with 10% FBS, 1% Penicillin/Streptomycin, L-glutamine and 50 µM β-mercaptoethanol (ME). γHV68 infected, and hygromycin selected A20.γHV68 (HE2.1) B cells [17] were obtained from Dr. Sam Speck (Emory University) and cultured in RPMI 1640 with 10% FBS, 1% Penicillin/Streptomycin + L-glutamine, 50 µM βME and 300 μg/mL Hygromycin B. A20 and A20.γHV68 B cells were treated with vehicle (untreated) or stimulated with 12-O-tetradecanolphrobol-13-actate (TPA) 20 ng/ml (Sigma) (in DMSO) harvested 24 hr later. BCBL-1 B cells, a body-cavity based B cell lymphoma cell line that is latently infected with KSHV (HHV-8), were obtained from the NIH AIDS reagent program (catalog # 3233). BCBL-1 cells were cultured in RPMI containing 20% FBS, 1% Penicillin/Streptomycin with L-glutamine, 1% HEPES and 50 µM βME. BCBL-1 B cells were treated with vehicle (untreated) or stimulated with 20 ng/ml TPA (in DMSO) and Sodium Butyrate (NaB) 0.3 mM (Calbiochem) (in water) and then harvested 72 hr later. BL41 B cells (negative for KSHV and EBV), were cultured in RPMI with 10% FBS, 1% Penicillin/Streptomycin with L-glutamine, and 50 µM βME. Mutu I cells, an EBV-infected, type I latency Burkitt’s lymphoma cell line [23] were obtained from Dr. Shannon Kenney (University of Wisconsin), and cultured in RPMI with 10% FBS, 1% Penicillin/Streptomycin and L-glutamine. Mutu I or BL41 B cells were either treated with vehicle (DMSO) or stimulated with 20 ng/ml TPA (in DMSO) and then harvested 48 hr later. EBV-immortalized B-lymphoblastoid cell lines (LCLs) were cultured in RPMI, 10% FBS, 1% Penicillin/Streptomycin with L-glutamine. LCL3, LCL9 and LCL209 BM were generated from Kenyan samples as previously described [35, 36].

### EBV infection of primary human B cells

Peripheral blood was obtained from consenting healthy adult donors and layered over Ficoll-Paque to isolate peripheral blood mononuclear cells (PBMCs). B cells were isolated from PBMCs through negative enrichment using EasySep Human B cell isolation kit (Stem Cell Technologies) following manufacturer’s protocol. B cells were plated at 1×10^6^ cells per mL in RPMI, 10% FBS, 1% Penicillin/Streptomycin with L-glutamine, then infected with EBV at a MOI of 10 genome copies per cell or mock infected. 5 days post infection cells were harvested for flow cytometry analysis. EBV virus stocks were generated from the EBV+ cell line B95.8 which was reactivated with TPA (50:50 EtOH:Acetone) and Sodium Butyrate (NaB) for 5 days. Supernatant was collected and centrifuged at 4,000xg for 10 min then passed over a 0.7 micron filter. The supernatant was then ultra-centrifuged at 16,000x g for 90 mins and resuspended in 1/200^th^ the initial volume using RPMI, 10% FBS, 1% Penicillin/Streptomycin with L-glutamine. Viral stocks were quantified following DNase treatment with qPCR analysis of the EBV BALF 5 gene done as previously described [37].

### Flow cytometric analysis

Cells were harvested at the indicated time points and processed for flow cytometry using the PrimeFlow™ RNA Assay (Thermo Fisher). Mouse cells were incubated with an Fc receptor blocking antibody (2.4G2) for 10 min and then fixed with 2% PFA (Fisher), washed with PBS (Life Technology). Cells were stained with a rabbit antibody against the γHV68 ORF4 protein, regulator of complement activation (RCA) [19], labeled with Zenon R-phycoerythrin rabbit IgG label reagent (Life Technologies) following manufacturer’s protocol. Human B cells were incubated in human Fc receptor blocking antibody then stained with Zombie Aqua fixable viability dye (1:500 dilution) (BioLegend), according to manufacturer’s protocol. Primary human B cells were subsequently stained with anti-CD19-FITC (Clone HIB19, dilution 1:25), and anti-CD69-PE antibody (clone FN50, dilution 1:20). Samples were subjected to the PrimeFlow™ RNA Assay following manufacturer’s protocols, using viral and host target probes conjugated to fluorescent molecules (Table S1). DAPI (BioLegend) was used on a subset of samples following manufacturer’s protocol, prior to PrimeFlow™ probe hybridization. Flow cytometric analysis was done on LSR II (BD Biosciences), Fortessa (BD Biosciences), and ZE5 (Bio-Rad) flow cytometers, with compensation values based on antibody-stained beads (BD Biosciences) and cross-validated using cell samples stained with individual antibody conjugates, with compensation modified as needed post-collection using FlowJo.

### Flow cytometric analysis utilizing barcoding

Cells were harvested at the indicated time points and processed for flow cytometry using the PrimeFlow™ RNA Assay (Thermo Fisher). Cells were incubated with an Fc receptor blocking antibody (2.4G2), then fixed and permeabilized using reagents from the PrimeFlow™ RNA Assay (Fixation Buffer 1 and Permeabilization Buffer). Samples from different experimental conditions were fluorescently barcoded, with cells treated with either: no fluorescent dye, Ghost Dye™ Violet 450 or Ghost Dye™ Violet 510 (Tonbo Biosciences, 1:200 dilution) according to manufacturer’s protocol. After washing, the three labeled samples were pooled together into a single sample and stained with Zenon-labeled polyclonal rabbit antisera against viral RCA as desribed above. Cells were fixed to cross-link antibody stain with Fixation Buffer 2 from PrimeFlow™ RNA Assay, then subjected to the PrimeFlow™ RNA Assay following manufacturer’s protocols. For the analysis of barcoded samples, singlet cells from barcoded samples were analyzed for fluorescence of either Ghost Dye™ Violet 450 or Ghost Dye™ Violet 510, to identify the three input populations: cells negative for Ghost 450 and Ghost 510, cells singly positive for Ghost 450, and cell singly positive for Ghost 510. To ensure that no artifacts were introduced due to assignment of one barcode to a specific experimental condition, barcodes used for each experimental condition were shuffled across independent samples.

### Imaging flow cytometry

Cells were treated as described above then harvested, and split into two aliquots: one for conventional flow cytometry, and one for imaging flow cytometry, acquired on an Amnis ImageStream®^X^ Mark II imaging flow cytometer (MilliporeSigma) with a 60X objective and low flow rate/high sensitivity using INSPIRE^®^ software. Brightfield (BF) and side scatter (SSC) images were illuminated by LED light and a 785nm laser respectively. Fluorescent probes were excited off 405nm, 488nm, and 642nm lasers with the power adjusted properly to avoid intensity saturation of the camera. Single color controls for compensation were acquired by keeping the same acquisition setting for samples, with the difference of turning the BF LED light and 785nm (SSC) laser off.

The acquired data were analyzed using IDEAS^®^ software (MilliporeSigma). Single cells that were in focus were defined as a population with a high “gradient RMS” value, an intermediate “Area” value, and a medium to high “Aspect ratio” value for subsequent analysis. Positive and negative events for each fluorescent marker were determined using the “Intensity” feature. TMER nuclear localization was quantified using “Similarity” feature, the log-transformed Pearson’s correlation coefficient by analyzing the pixel values of two image pairs [27]. The degree of nuclear localization of TMER was measured by correlating the pixel intensity of two images with the same spatial registry. The paired TMER and DAPI images were quantified by measuring the “Similarity Score” which cells with high similarity scores display high TMER nuclear localization with similar image pairs. By contrast cells with low similarity scores show low TMER nuclear localization with dissimilar image pairs. Cell cycle stage analysis was performed on data obtained from the imaging flow cytometer, stratifying cells based on DAPI content to identify cells in the G0/G1 phase, S phase or G2/M phase of the cell cycle.

### Cell purification and DNA quantitation based on viral gene expression

Mouse fibroblasts (3T12) were infected with WT γHV68 (MOI=5), harvested and processed for PrimeFlow™ at 16 hpi, followed by cell sorting using a BD FACSAria to purify cells based on relative TMER and ORF73 expression. Post-sort purity checks on sorted populations indicated that cell purities were 100% for TMER-ORF73- cells, 94.1% for TMER+ ORF73- and 98% for TMER+ORF73+ cells. DNA was isolated from samples using the DNeasy Blood and Tissue Kit (Qiagen), with an overnight proteinase K incubation followed by heat inactivation (95°C for 10 min). DNA was precipitated using ammonium acetate alcohol. 40 ng of DNA per sample was subjected to qPCR analysis using LightCycler 480 Probe Master-Mix kit (Roche) and primer sets for γHV68 gB and host NFAT5 (Table S3). gB standard curve was generated using a gB plasmid dilution series ranging from 10^10^ to 10^1^ copies diluted in background DNA, with a limit of detection (LOD) of 100 copies [38]. Host NFAT5 standard curve was generated using HEK293 cells, with 2×10^5^ cell equivalents subjected to serial 10-fold dilutions in background, salmon sperm DNA, from 10^5^ to 10^1^ copies (LOD=10 copies). Quantitation of viral gB copy number was standardized relative to input material using the formula (gB copy number / NFAT5 copy number) / 2, based on the assumption that cells have 2 copies of NFAT5 gene.

### KSHV genome quantification

BCBL-1 or BL41 cells were plated at 7.5e5 cells/well in a 6 well plate with 20 ng/ml TPA and 0.3 mM NaB or vehicle only (DMSO and H_2_O). Cells and supernatant were harvested at 72 hrs post-treatment, hard-spun for 30 min at 4° C and DNA was isolated using the DNeasy Blood and Tissue kit following manufacturer’s protocol, except for sample digestion for 1 hour instead of 10 min. 100 ng of DNA per sample was used for qPCR analysis via SYBR green detection using KSHV ORF50 primers (5’ -TCC GGC GGA TAT ACC GTC AC- 3’ and 5’- GGT GCA GCT GGT ACA GTG TG-3’) [39]. qPCR was analyzed using relative quantification normalized against unit mass calculation, ratio = E^deltaCt^ (Real-Time PCR Application Guide, Bio-Rad Laboratories Inc. 2006).

### RNA and qRT-PCR

RNA was isolated using Trizol (Life Technologies) per manufacturer’s protocol and re-suspended in DEPC treated water. 3 μg of RNA was treated with DNase 1 (Promega) for 2 hours at 37°C, heat inactivated for 10 min at 65°C. 500 ng of RNA was then subjected to reverse transcription using SuperScript II (Life Technologies) following manufacturer’s protocol for gene specific, oligo(dT), or random primers (Life Technologies). Quantitative PCR (qPCR) was performed using iQ SYBR Green super mix (Bio-Rad) follow manufacturer’s protocol using host and viral primer sets (Table S2) or using QuantiTech Primer Assay (Qiagen) for 18s (Hs-RRN18S_1_SG). qPCR conditions: 3 min at 95°C, amplification cycles for 40 cycles of 15 sec at 95°C, annealing/ extension at temperature for specific primer set for 1 min ending with a melt curve which started at 50°C or 55°C to 95°C increasing 0.5°C for 0:05 sec. A standard curve for each primer set was generated by pooling a portion of each sample together and doing a 1:3 serial dilution. 75 ng of cDNA of the unknown samples was loaded per qPCR reaction/primer set, with reactions run on a Bio-Rad 384 CFX LightCycler and data analyzed using Bio-Rad CFX manager software. Data analysis was done using the 1:3 standard curve as the control Ct value to calculate the delta ct, and the Pfaffl equation was used to define the fold difference between the gene of interest and 18s (reference gene) [40]. qPCR products were analyzed by melt curve analysis, with all reactions having a prominent, uniform product. In the case of primers with an aberrant melt curve product (e.g. that arose at late cycles), products were clearly a different product as defined by melt curve analysis.

### Software and Statistical analysis

All flow cytometry data were analyzed in FlowJo (version 8.8.7, 10.5.0, and 10.5.3), with flow cytometry data shown either as histogram overlays or pseudo-color dot plots (with or without smoothing), showing outliers (low or high resolution) on log_10_ scales. Statistical analysis and graphing were done in GraphPad Prism (Version 6.0d and 7.0d). Statistical significance was tested by unpaired t test (when comparing two conditions) or by one-way ANOVA (when comparing three or more samples) subjected to multiple corrections tests using recommended settings in Prism. *<u>X-shift analysis:</u>* For automated mapping of flow cytometry data using X-shift, data were obtained from compensated flow cytometry files, exported from FlowJo, using singlets that were live (defined by sequential gating on single cells by FSC-H vs. FSC-W and SSC-H vs. SSC- W, that were DAPI bright vs. SSC-A). These events were imported into the Java based program VorteX (http://web.stanford.edu/~samusik/vortex/) [26]. Four parameters [TMER (AlexaFluor (AF) 488), RCA (PE), ORF73 (AF647), and Actin (AF750)] were selected for clustering analysis using the X-shift algorithm. The following settings were used when importing the data set into VorteX: i) Numerical transformation: arcsinh(x/f), f=150, ii) noise threshold: apply noise threshold of 1.0 (automatic and recommended setting), iii) feature rescaling: none, and iv) normalization: none, v) a Euclidean noise filter was used with a Minimal Euclidean length of the profile of 1.0, and vi) an import max of 1,000 rows from each file after filtering was selected. The following settings were used when preparing the data set for clustering analysis: i) distance measure: angular distance, ii) clustering algorithm: X-shift (gradient assignment), iii) density estimate: N nearest neighbors (fast), iv) number of neighbors for density estimate (K): from 150 to 5, with 30 steps, and v) number of neighbors for mode finding (N): determine automatically. After the cluster analysis was completed, all results were selected and the K value that corresponded with optimal clustering (the elbow point) was calculated, in this case K= 50. All clusters (seven clusters total) for the optimal K value were selected and a Force-Directed Layout was created. The maximum number of events sampled from each cluster was 20, and the number of nearest neighbors was 10. All settings used for this analysis were automated or explicitly recommended (https://github.com/nolanlab/vortex/wiki). Force-Directed layouts in Fig. 6A were saved as graphml files from VorteX, opened in the application Gephi v 0.9.1, and colored by different variables (Cluster ID, experimental group, Actin mRNA, RCA, ORF73, and TMERs respectively) in Adobe Illustrator CC 2017. Full details on use of the X-shfit algorithm and analysis pipeline can be found in [41]. *<u>tSNE analysis:</u>* Gated events for each of the six identified populations were exported from FlowJo, and then imported into Cytobank (www.cytobank.org) for analysis using the viSNE algorithm. Each file was used for a separate viSNE analysis (six total runs), where all available events were selected for clustering (202,669, 128,028, 29,096, 5,610, 8,956, 35,850 respectively) and four parameters were selected for clustering (Actin mRNA, RCA, ORF73, and TMERs). The resulting tSNE plots were colored according to expression using the “rainbow” color option, with individual events shown using the stacked dot option. The channel range was user-defined for each marker according to the range in expression established in Fig. 6E.

## ACKNOWLEDGMENTS

The authors would like to acknowledge insightful comments made by members of the Clambey and van Dyk laboratories, technical assistance by Eva Medina, pilot studies establishing the feasibility of fluorescent barcoding by Shannon Rudolph, support of flow cytometry services through ClinImmune, the Dept. of Immunology & Microbiology, and the University of Colorado Cancer Center, and expert technical guidance from Matt Cato and Dr. Nori Ueno (Thermo Fisher). T.C. and B.A. are employees of EMD Millipore and Luminex respectively and have a potential conflict of interest. E.T.C. was a recipient of the 2016 North America Affymetrix Single-cell Grant Recipient, for studies unrelated to this manuscript.

## AUTHOR CONTRIBUTIONS

<u>Conceptualization:</u> Eric T. Clambey, Linda F. van Dyk

<u>Data curation:</u> Lauren M. Oko, Eric T. Clambey

<u>Formal analysis:</u> Lauren M. Oko, Abigail K. Kimball, Rachael E. Kaspar, Benjamin Alderete, Tim Chang, Eric T. Clambey

<u>Funding acquisition:</u> Eric T. Clambey, Linda F. van Dyk

<u>Investigation:</u> Lauren M. Oko, Ashley N. Knox, Carrie B. Coleman

<u>Methodology:</u> Lauren M. Oko, Carrie B. Coleman, Rosemary Rochford, Benjamin Alderete, Tim Chang, Linda F. van Dyk, Eric T. Clambey

<u>Project administration:</u> Eric T. Clambey, Linda F. van Dyk

<u>Resources:</u> Lauren M. Oko, Rosemary Rochford, Benjamin Alderete, Tim Chang

<u>Software:</u> Abigail K. Kimball

<u>Supervision:</u> Eric T. Clambey, Linda F. van Dyk

<u>Validation:</u> Lauren M. Oko, Eric T. Clambey

<u>Visualization:</u> Lauren M. Oko, Abigail K. Kimball, Rachael E. Kaspar, Benjamin Alderete, Tim Chang, Eric T. Clambey

<u>Writing – original draft:</u> Eric T. Clambey, Linda F. van Dyk

<u>Writing – review & editing:</u> Lauren M. Oko, Linda F. van Dyk, Eric T. Clambey

## SUPPORTING INFORMATION LEGENDS

**Supplemental Table 1. Viral and host probes for PrimeFlow™ analysis.**

**Supplemental Table 2. qPCR primer sets.**

**Supplemental Table 3. TaqMan PCR probe sets used in this study.**

**Supplemental Figure 1. Analysis of KSHV genomic DNA in BCBL-1 cells either untreated or stimulated.** Data depict qPCR analysis of KSHV genomic DNA, quantified using PCR primers for the KSHV gene 50. Fold change in KSHV DNA was calculated relative to untreated BCBL-1 cells, using ΔCt normalized to mass unit, defined using 100 ng DNA. Data compiled from 3 independent experiments for BCBL-1 cultures, with 3 biological replicates per experiment. Data from BL-41 cells derived from a single experiment, with 3 biological replicates. Data depict mean +/- SEM, with individual biological replicates depicted using individual symbols. Statistical analysis of the difference between untreated and stimulated samples was done using an unpaired t test, with Welch’s correction, with statistical significance defined by ***, p<0.001.

**Supplemental Figure 2. Analysis of EBER expression by PrimeFlow™ in a panel of human B cell lines.** (A) Comparison of fluorescence in BL41 (EBV-) and Mutu I (EBV+) cells, where cell lines were either incubated with no probe or an EBER probe. The three populations identified on these plots, indicated by 3 polygons with black lines, correspond to: left, the true negative population; middle, background fluorescence observed in BL41 cells stained with EBER probes; right, EBER+ events defined by expression above these two different thresholds. (B) Quantitation of the frequency of EBER+ events among viable cells across all the cell lines analyzed, with symbols depicting individual replicate data, bars showing mean ± SEM. (C) Analysis of EBER expression in three different LCL cultures, as indicated, comparing background fluorescence (No probe) with fluorescence following EBER probe hybridization. (D) Gating hierarchy to analyze EBER fluorescence in viable cells. All flow cytometry plots show events defined as lymphocytes by forward and side scatter, doublet discrimination, and viable cells defined by exclusion of a viability dye. Data are from a single experiment with one to three biological replicates (n=1 Mutu I, n=3 for BL41 and LCLs).

**Supplemental Figure 3. Analysis of EBER expression by PrimeFlow™ in human primary B cells subjected to in vitro EBV infection.** (A) Comparison of EBER expression in human primary B cells subjected to either mock or EBV infection (10 genome copies/cell) for 5 days. Cells were either incubated with a cocktail of antibodies and the EBER probe, or with specific exclusion of the EBER probe (i.e. a full minus one control, No probe). Data depict the frequency of EBER+ events within lymphocytes that were singlets and CD19+ B cells. Data are from a single experiment. (B) Comparison of EBER expression in human primary B cells subjected to either mock or EBV infection (10 genome copies/cell) for 5 days. Data depict the frequency of EBER+ events within lymphocytes that were viable, singlets, and CD19+ B cells. (C) Quantitation of the frequency of EBER+ cells among viable B cells in mock or EBV infected cultures, with symbols depicting individual replicate data, bars showing mean ± SEM. (D) Comparison of cell characteristics in EBV infected cells, between EBER- (in black) and EBER+ (in red) events using histogram overlays, with populations defined in panel B. Each histogram overlay depicts three biological replicates, comparing expression of the defined parameter between EBER- and EBER+ populations.

**Supplemental Figure 4. Measurement of viral gene expression heterogeneity by multiple measures.** 3T12 cells were infected with WT γHV68 at an MOI=5 and analyzed at 16 hpi. **(A)** Histogram overlays comparing expression of the indicated parameters between 5 different populations of cells stratified by TMER and ORF73 expression, as defined on the leftmost panel, with data representative of three biological replicates. **(B)** Histogram overlays comparing expression of the indicated parameter in mock (black) or WT γHV68 -infected (red) samples, comparing three biological replicates with events defined as DAPI+ singlets. **(C)** Viral DNA as a function of viral gene expression defined by TMER and 73 expression. NIH 3T12 cells were either mock or virus infected (MOI=5), harvested at 16 hpi, collected in bulk (mock and infected, no sort) or subject to FACS purification based on expression of TMER and 73 expression to isolate TMER- 73-, TMER+ 73- and TMER high 73 high populations. DNA was harvested from all samples, subjected to qPCR with primers to the gB gene of gHV68. DNA content was normalized relative to cellular DNA, defined by parallel amplification of the NFAT5 gene. Data depict mean ± SEM, with symbols indicating individual samples. Statistical analysis was performed using one-way ANOVA subjected to Tukey’s multiple comparisons test, comparing viral DNA content between sorted cell populations, with statistical significance as identified, *** p<0.001. **(D)** Viral gene expression during primary lytic replication with gHV68 is relatively comparable between cells in different stages of the cell cycle stages. Data are from imaging flow cytometry-based studies, depicting the frequency of cells stratified by expression of TMERs and ORF73 (as defined in Fig. 6D) present among total DAPI+ cells (top row) compared to cells in either the G0/G1, S or G2/M phase of the cell cycle. Data depict results from three independent experiments with 8 total replicates, with columns plotting mean ± SEM. Flow cytometry data depict single cells, defined by sequential removal of doublets according to SSC-A x SSC-H and FSC-A x FSC-H.

**Supplemental Figure 5. tSNE analysis of cell size and granularity as a function of infection and gene expression status.** Data show flow cytometry data using populations defined in Fig. 6. Data show all DNA+ (DAPI+) single cells (FSC-A, SSC-A) subjected to the tSNE dimensionality reduction algorithm, depicting relative expression values for cell size (FSC) and granularity (SSC) in rows relative to the defined cell populations (columns). The tSNE algorithm provides each cell with a unique coordinate, displayed on a two-dimensional plot (tSNE1 versus tSNE2), such that FSC and SSC values within cellular islands can be directly compared to the corresponding cell islands presented in Fig. 7C. The channel range was locally-defined for each individual and channel via Cytobank. Flow cytometry data shows single cells that are DNA+ (DAPI+). Data are from three independent experiments.

**Supplemental Figure 6. Phosphonoacetic acid treatment alters viral gene expression during lytic replication.** 3T12 fibroblasts were infected with WT gHV68 (MOI=5), either in the absence of phosphonoacetic acid (no PAA) or incubated with PAA (200 mg/mL, +PAA), harvested at 18 hpi and subjected to PrimeFlow analysis. (A,B) Analysis of TMER and ORF73 expression profiles on a biaxial plot (A), with total data shown in (B). (C,D) Analysis of TMER and Actin mRNA expression profiles on a biaxial plot (C), with total data shown in (D). Events were gated on cells subjected to doublet discrimination. Data depict mean ± SEM from two independent experiments, with 4 biological replicates per group denoted by individual symbols. Statistical analysis done using either a two-way ANOVA with Sidak’s multiple correction test (panel B) or an unpaired t test (panel D), with statistical significance as identified ****, p<0.0001.

