## Supplemental Figures 1-6 for "Multidimensional analysis of Gammaherpesvirus RNA expression reveals unexpected heterogeneity of gene expression"

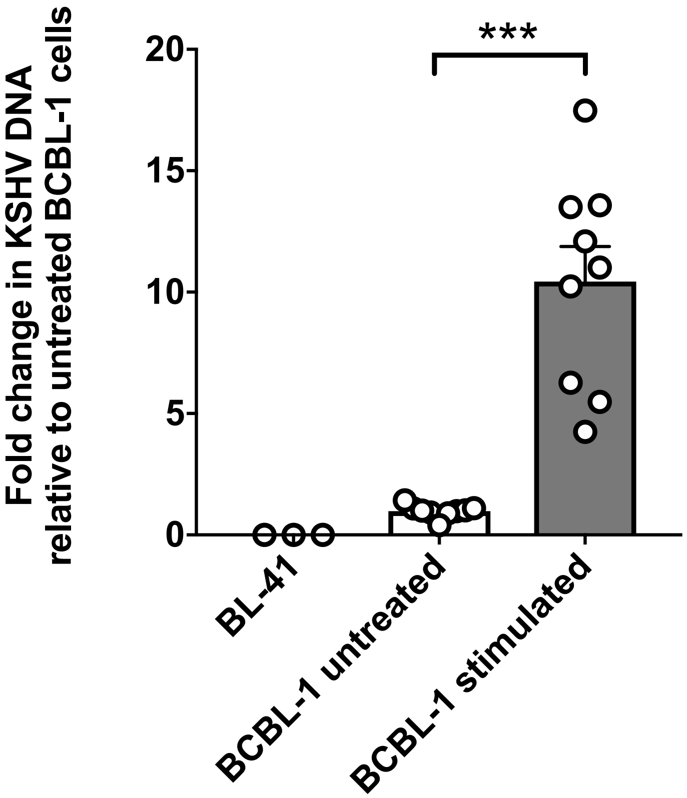


**Supplemental Figure 1. Analysis of KSHV genomic DNA in BCBL-1 cells either untreated or stimulated.** Data depict qPCR analysis of KSHV genomic DNA, quantified using PCR primers for the KSHV gene 50. Fold change in KSHV DNA was calculated relative to untreated BCBL-1 cells, using ΔCt normalized to mass unit, defined using 100 ng DNA. Data compiled from 3 independent experiments for BCBL-1 cultures, with 3 biological replicates per experiment. Data from BL-41 cells derived from a single experiment, with 3 biological replicates. Data depict mean +/- SEM, with individual biological replicates depicted using individual symbols. Statistical analysis of the difference between untreated and stimulated samples was done using an unpaired t test, with Welch's correction, with statistical significance defined by ***, $p<0.001$.

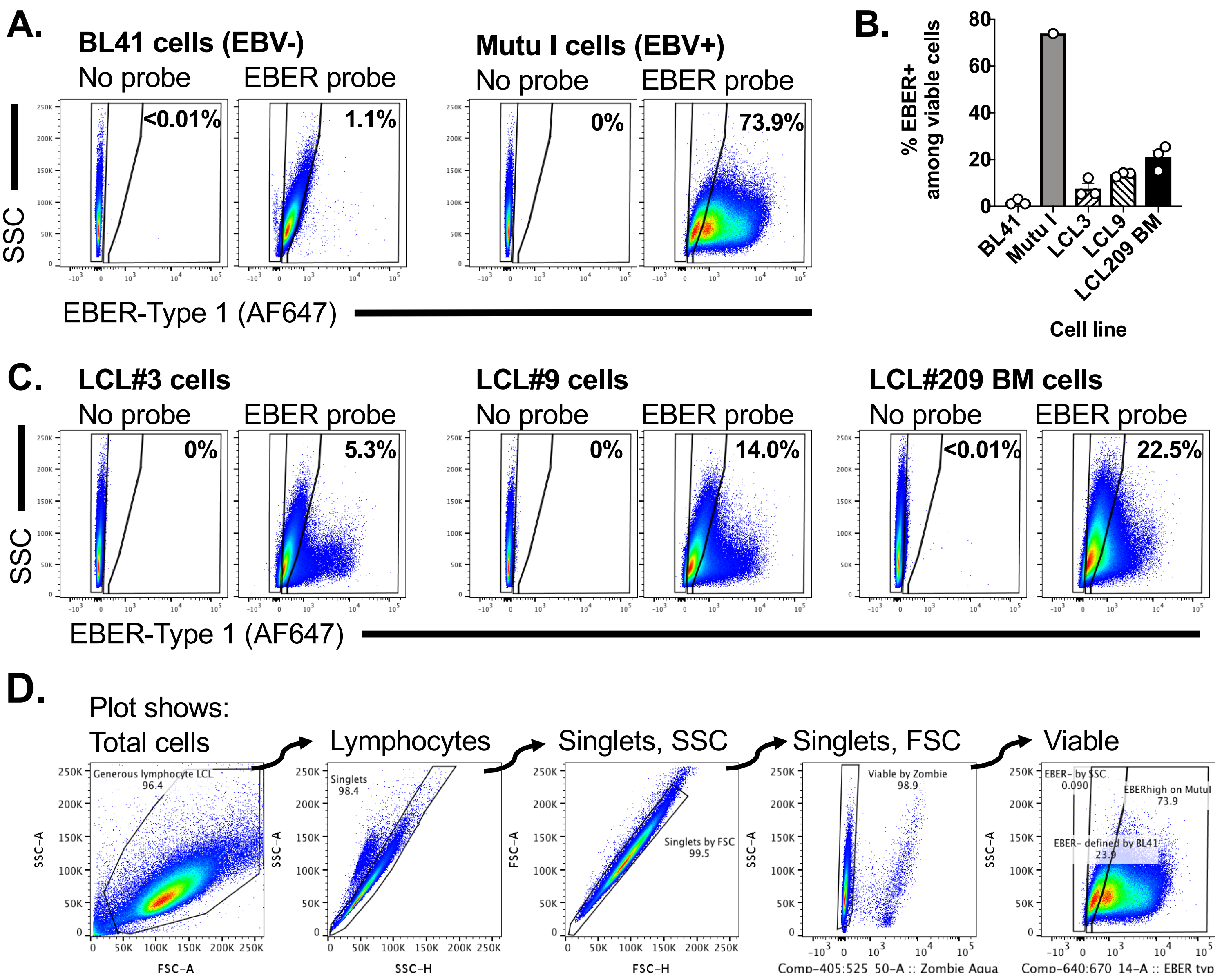


**Supplemental Figure 2. Analysis of EBER expression by PrimeFlow™ in a panel of human B cell lines.** (A) Comparison of fluorescence in BL41 (EBV-) and Mutu I (EBV+) cells, where cell lines were either incubated with no probe or an EBER probe. The three populations identified on these plots, identified by 3 polygons with black lines, correspond to: left, the true negative population; middle, background fluorescence observed in BL41 cells stained with EBER probes; right, EBER+ events defined by expression above these two different thresholds. (B) Quantitation of the frequency of EBER+ events among viable cells across all the cell lines analyzed, with symbols depicting individual replicate data, bars showing mean ± SEM. (C) Analysis of EBER expression in three different LCL cultures, as indicated, comparing background fluorescence (No probe) with fluorescence following EBER probe hybridization. (D) Gating hierarchy to analyze EBER fluorescence in viable cells. All flow cytometry plots show events defined as lymphocytes by forward and side scatter, doublet discrimination, and viable cells defined by exclusion of a viability dye. Data are from a single experiment with one to three biological replicates (n=1 Mutu I, n=3 for BL41 and LCLs).

Oko et al
Supplemental Figure 2

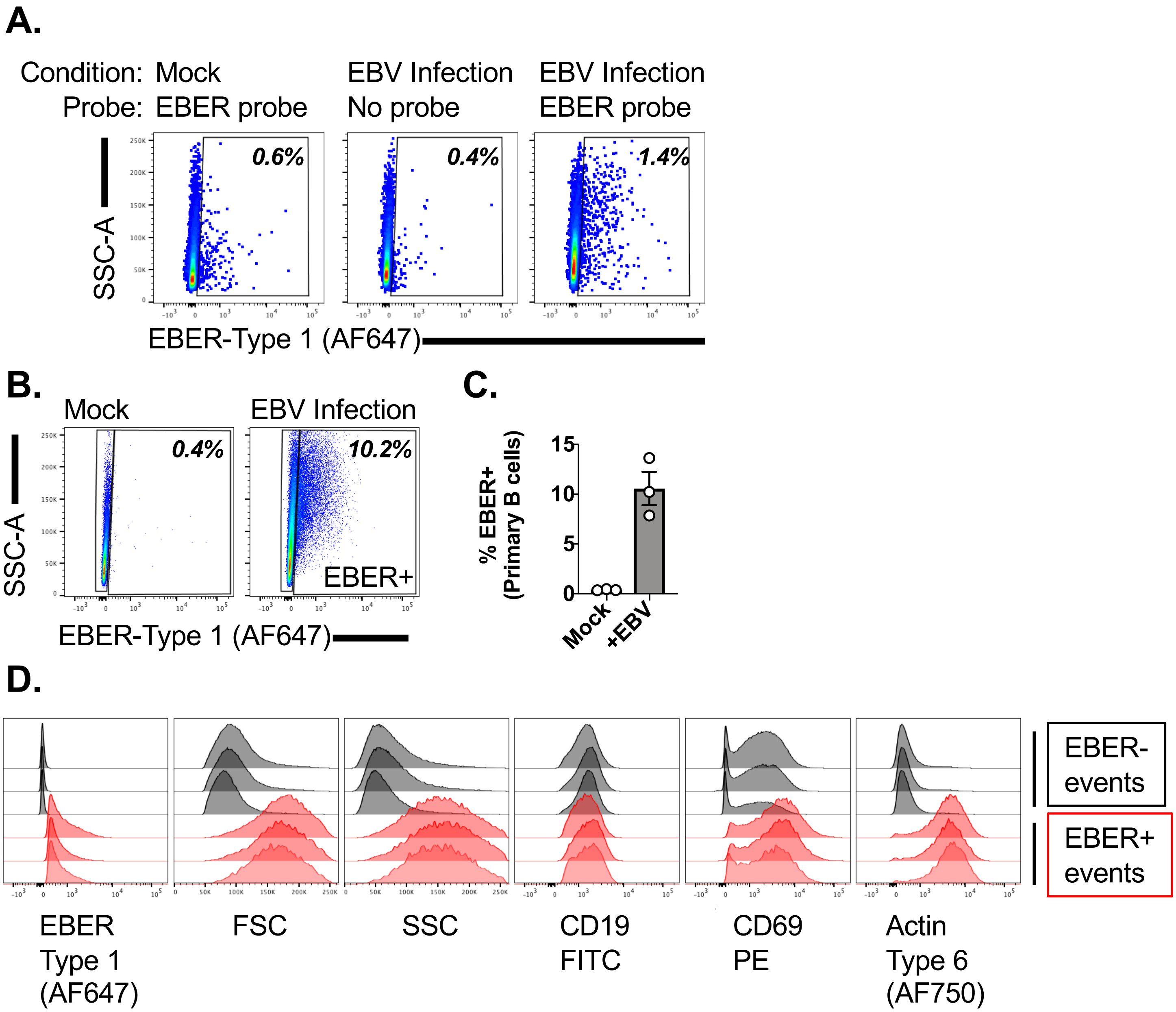


**Supplemental Figure 3. Analysis of EBER expression by PrimeFlow™ in human primary B cells subjected to in vitro EBV infection.** (A) Comparison of EBER expression in human primary B cells subjected to either mock or EBV infection (10 genome copies/cell) for 5 days. Cells were either incubated with a cocktail of antibodies and the EBER probe, or with specific exclusion of the EBER probe (i.e. a full minus one control, No probe). Data depict the frequency of EBER+ events within lymphocytes that were singlets and CD19+ B cells. Data are from a single experiment. (B) Comparison of EBER expression in human primary B cells subjected to either mock or EBV infection (10 genome copies/cell) for 5 days. Data depict the frequency of EBER+ events within lymphocytes that were viable, singlets, and CD19+ B cells. (C) Quantitation of the frequency of EBER+ cells among viable B cells in mock or EBV infected cultures, with symbols depicting individual replicate data, bars showing mean ± SEM. (D) Comparison of cell characteristics in EBV infected cells, between EBER- (in black) and EBER+ (in red) events using histogram overlays, with populations defined in panel B. Each histogram overlay depicts three biological replicates, comparing expression of the defined parameter between EBER- and EBER+ populations.

Oko et al
Supplemental Figure 3

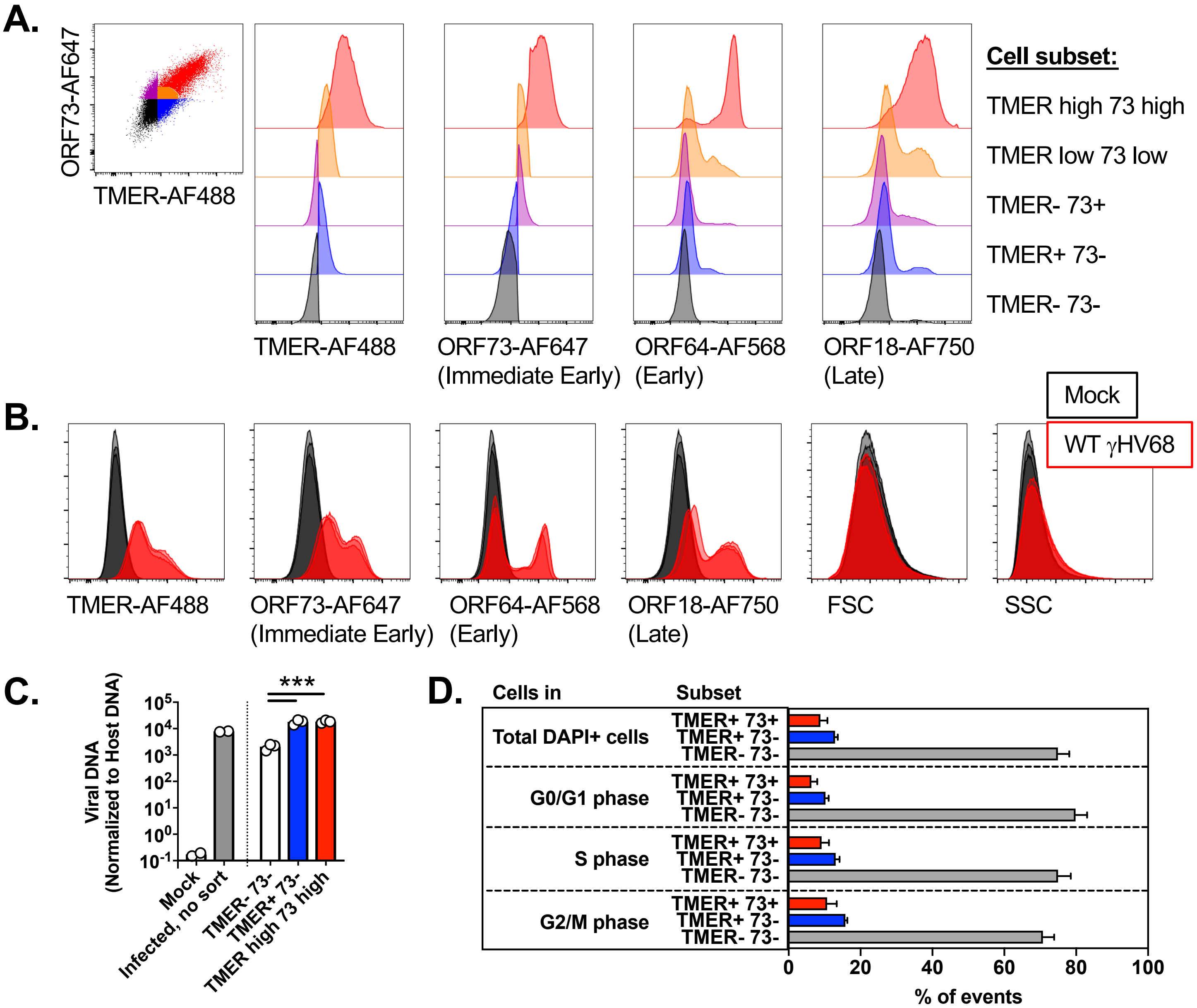


**Supplemental Figure 4. Measurement of viral gene expression heterogeneity by multiple measures.** 3T12 cells were infected with WT γHV68 at an MOI=5 and analyzed at 16 hpi. **(A)** Histogram overlays comparing expression of the indicated parameters between 5 different populations of cells stratified by TMER and ORF73 expression, as defined on the leftmost panel, with data representative of three biological replicates. **(B)** Histogram overlays comparing expression of the indicated parameter in mock (black) or WT γHV68-infected (red) samples, comparing three biological replicates with events defined as DAPI+ singlets. **(C)** Viral DNA as a function of viral gene expression defined by TMER and 73 expression. NIH 3T12 cells were either mock or virus infected (MOI=5), harvested at 16 hpi, collected in bulk (mock and infected, no sort) or subject to FACS purification based on expression of TMER and 73 expression to isolate TMER- 73-, TMER+ 73- and TMER high 73 high populations. DNA was harvested from all samples, subjected to qPCR with primers to the gB gene of γHV68. DNA content was normalized relative to cellular DNA, defined by parallel amplification of the NFAT5 gene. Data depict mean ± SEM, with symbols indicating individual samples. Statistical analysis was performed using one-way ANOVA subjected to Tukey's multiple comparisons test, comparing viral DNA content between sorted cell populations, with statistical significance as identified, *** $p<0.001$. **(D)** Viral gene expression during primary lytic replication with γHV68 is relatively comparable between cells in different stages of the cell cycle stages. Data are from imaging flow cytometry-based studies, depicting the frequency of cells stratified by expression of TMERs and ORF73 (as defined in Fig. 6D) present among total DAPI+ cells (top row) compared to cells in either the G0/G1, S or G2/M phase of the cell cycle. Data depict results from three independent experiments with 8 total replicates, with columns plotting mean ± SEM. Flow cytometry data depict data from single cells, defined by sequential removal of doublets according to SSC-A x SSC-H and FSC-A x FSC-H.

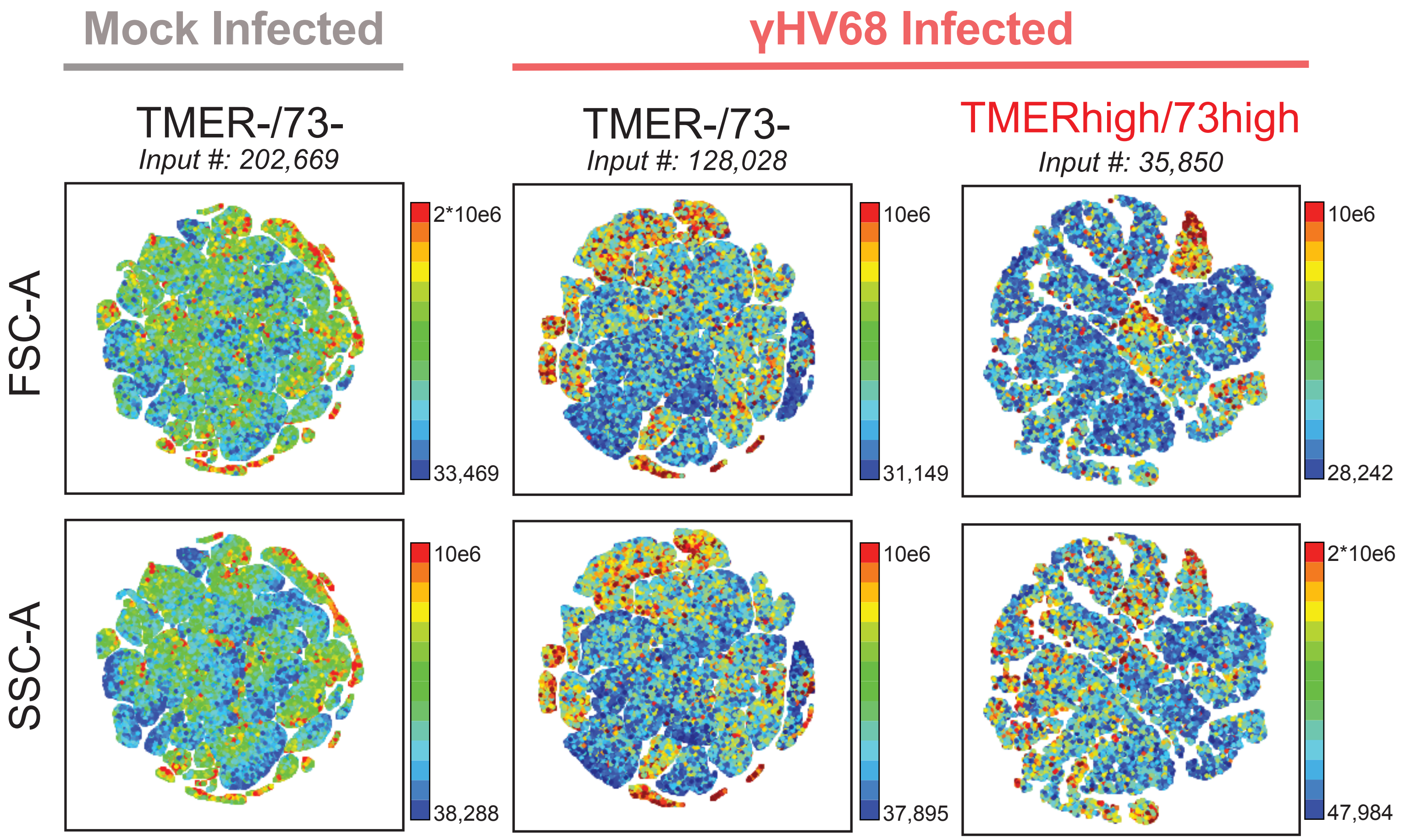


**Supplemental Figure 5. tSNE analysis of cell size and granularity as a function of infection and gene expression status.** Data show flow cytometry data using populations defined in Fig. 6. Data show all DNA+ (DAPI+) single cells (FSC-A, SSC-A) subjected to the tSNE dimensionality reduction algorithm, depicting relative expression values for cell size (FSC) and granularity (SSC) in rows relative to the defined cell populations (columns). The tSNE algorithm provides each cell with a unique coordinate, displayed on a two-dimensional plot (tSNE1 versus tSNE2), such that FSC and SSC values within cellular islands can be directly compared to the corresponding cell islands presented in Fig. 7C.
The channel range was locally-defined for each individual and channel via Cytobank. Flow cytometry data shows single cells that are DNA+ (DAPI+). Data are from three independent experiments.

**Supplemental Figure 5.**
**Oko et al**

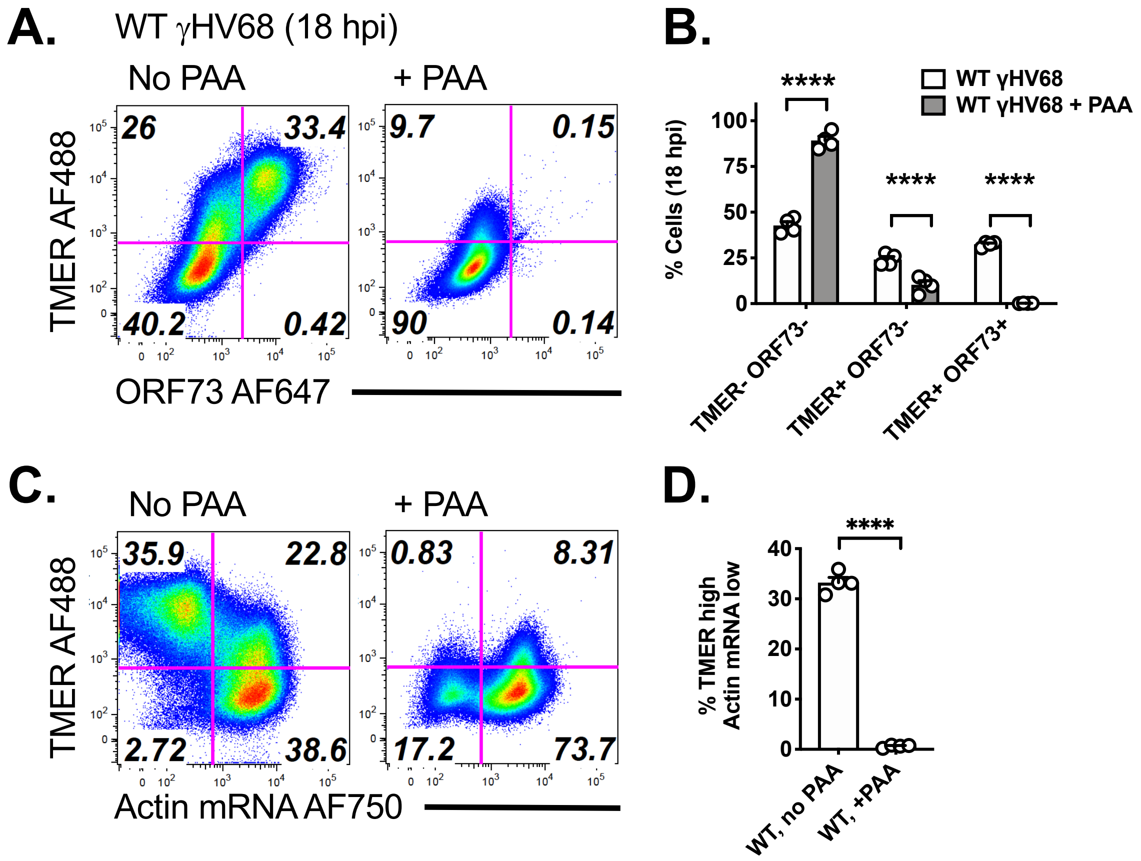


**Supplemental Figure 6. Phosphonoacetic acid treatment alters viral gene expression during lytic replication.** 3T12 fibroblasts were infected with WT γHV68 (MOI=5), either in the absence of phosphonoacetic acid (no PAA) or incubated with PAA (200 μg/mL, +PAA), harvested at 18 hpi and subjected to PrimeFlow analysis. (A,B) Analysis of TMER and ORF73 expression profiles on a biaxial plot (A), with total data shown in (B). (C,D) Analysis of TMER and Actin mRNA expression profiles on a biaxial plot (C), with total data shown in (D). Events were gated on cells subjected to doublet discrimination. Data depict mean ± SEM from two independent experiments, with 4 biological replicates per group denoted by individual symbols. Statistical analysis done using either a two-way ANOVA with Sidak's multiple correction test (panel B) or an unpaired t test (panel D), with statistical significance as identified ****, $p<0.0001$.
