## Supplemental Tables 1-3 for "Multidimensional analysis of Gammaherpesvirus RNA expression reveals unexpected heterogeneity of gene expression"

### Supplemental Data Tables for "Multidimensional analysis of Gammaherpesvirus RNA expression reveals unexpected heterogeneity of gene expression"

Lauren M. Oko[1], Abigail K. Kimball[2], Rachael E. Kaspar[2], Ashley N. Knox[1], Carrie B. Coleman[1], Rosemary Rochford[1], Tim Chang[3], Benjamin Alderete[4], Linda F. van Dyk[1]*, Eric T. Clambey[2]*#

**Affiliations:**

[1] Department of Immunology and Microbiology, [2] Department of Anesthesiology, University of Colorado Denver | Anschutz Medical Campus, Aurora, CO, 80045, USA

[3] MilliporeSigma, a business of Merck KGaA, Darmstadt, Germany (Seattle, WA)

[4] Luminex Corporation, Austin, TX, USA

*** Co-Corresponding authors:**

[#] **Lead author** for MS correspondence

**Short title:** Single-cell heterogeneity of Gammaherpesvirus RNA expression

Supplemental Table 1
Oko et al

Table S1: Viral and host probes for PrimeFlow™ analysis

| **Target and probe name** | **Fluorochrome** | **ThermoFisher Probe number** | **Target species** |
|---|---|---|---|
| Murid Herpesvirus ORF18 Type 4 | Alexa Fluor® 488 | VF4-6000512 | γHV68 |
| Murid Herpesvirus ORF18 Type 6 | Alexa Fluor® 750 | VF6-6001307 | γHV68 |
| Murid Herpesvirus ORF64 Type 10 | Alexa Fluor® 568 | VF10-6001306 | γHV68 |
| Murid Herpesvirus ORF72 Type 1 | Alexa Fluor® 647 | VF1-20941 | γHV68 |
| Murid Herpesvirus ORF73 Type 1 | Alexa Fluor® 647 | VF1-17077 | γHV68 |
| Murid Herpesvirus TMER Type 1 | Alexa Fluor® 647 | VF1-17076 | γHV68 |
| Murid Herpesvirus TMER Type 4 | Alexa Fluor® 488 | VF4-17586 | γHV68 |
| Mouse β-Actin Type 1 | Alexa Fluor® 647 | VB1-10350 | *Mus musculus* |
| Mouse β-Actin Type 4 | Alexa Fluor® 488 | VB4-10432 | *Mus musculus* |
| Mouse β-Actin Type 6 | Alexa Fluor® 750 | VB6-12823 | *Mus musculus* |
| HHV4 EBER 1-2 Type 1 | Alexa Fluor® 647 | VF1-12409 | EBV |
| HHV8 ORF73 Type 1 | Alexa Fluor® 647 | VF1-6000252 | KSHV |
| HHV8 T1.1 Type 4 | Alexa Fluor® 488 | VF4-6000059 | KSHV |
| Human β-Actin Type 6 | Alexa Fluor® 750 | VA6-10506 | *Homo sapiens* |
| Human IL-6 Type 10 | Alexa Fluor® 568 | VA10-13146 | *Homo sapiens* |

Supplemental Table 2
Oko et al

Table S2: qPCR primer sets used in this study

| **PCR primer** | **Sequence 5'-3'** | **Tm °C** |
|---|---|---|
| HV68 tRNA5 sense | GCC AGG GTA GCT CAA TTG | 52 °C |
| miR-M1-12 RLMRTPCR | AAG GGG TAG GAC TCC CAC | 52 °C |
| Mouse bActin Sense | GCC ACC AGT TCG CCA TGG | 56 °C |
| Mouse bActin Antisense Inner | CAG GGT CAG GAT ACC TCT CTT G | 56 °C |
| ORF73 RT-PCR Reverse Primer | GAG CCC CCT ACA GAG CCC CC | 59 °C |
| ORF73 RT-PCR Forward Primer | CAC CTT GCT CAC CGG CA | 59 °C |
| Hu bActin SYBR green Forward | GAT GAG ATT GGC ATG GCT TT | 60 °C |
| Hu bActin SYBR green Reverse | CAC CTT CAC CGT TCC AGT TT | 60 °C |
| KSHV ORF50 Forward | TCC GGC GGA TAT ACC GTC AC | 60 °C |
| KSHV ORF50 Reverse | GGT GCA GCT GGT ACA GTG TG | 60 °C |

Supplemental Table 3
Oko et al

Table S3: TaqMan PCR probe sets used in this study

| **Primer/Probe Name** | **Sequence (5’-3’)** | **Annealing Temp. (C°)** |
|---|---|---|
| γHV68 Forward Primer | GGC CCA AAT TCA ATT TGC CT | 60 C° |
| γHV68 Reverse Primer | CCC TGG ACA ACT CCT CAA GC | 60 C° |
| γHV68 Probe FAM/TAMRA | ACA AGC TGA CCA CCA GCG TCA ACA AC | 60 C° |
| Host NFAT5 Forward Primer | CAT GAG CAC CAG TTC CTA CAA TGAT | 60 C° |
| Host NFAT5 Reverse Primer | TGC TTT GGA TTT CGT TTT CGT GAT T | 60 C° |
| Host NFAT5 Probe VIC/HBQ | ACG AGG TAC CTC AGT GTT | 60 C° |
